# Unexpectedly High Subclonal Mutational Diversity in Human Colorectal Cancer and Its Significance

**DOI:** 10.1101/672493

**Authors:** Lawrence A. Loeb, Brendan F. Kohrn, Kaitlyn J. Loubet-Senear, Yasmin J. Dunn, Eun Hyun Ahn, Jacintha N. O’Sullivan, Jesse J. Salk, Mary P. Bronner, Robert A. Beckman

## Abstract

Human colorectal cancers (CRC) contain both clonal and subclonal mutations. Clonal mutations are positively selected, present in most cells and drive malignant progression. Subclonal mutations are randomly dispersed throughout the genome; they provide a vast reservoir of mutant cells that can expand, repopulate the tumor and result in the rapid emergence of resistance, as well as being a major contributor to tumor heterogeneity. Here, we apply Duplex Sequencing (DS) methodology to quantify subclonal mutations in CRC tumor with unprecedented depth (10^4^) and accuracy (<10^−7^). We measured mutation frequencies in genes encoding replicative DNA polymerases and in genes frequently mutated in CRC, and found an unexpectedly high effective mutation rate, 7.1 × 10^−7^. The curve of subclonal mutation accumulation as a function of sequencing depth, using DNA obtained from five different tumors, is in accord with a neutral model of tumor evolution. We present a new theoretical approach to model neutral evolution independent of the infinite sites assumption (which states that a particular mutation arises only in one tumor cell at any given time). Our analysis indicates that the infinite sites assumption is not applicable once the number of tumor cells exceeds the reciprocal of the mutation rate, a circumstance relevant to even the smallest clinically diagnosable tumor. Our methods allow accurate estimation of the total mutation burden in clinical cancers. Our results indicate that no DNA locus is wild type in every malignant cell within a tumor at the time of diagnosis (probability of all cells wild type = 10^−308^).

**Significance Statement:** Cancers evolve many mutations. Clonal mutations are selected early. Subsequent evolution occurs in a branching fashion, possibly without selection (“neutral evolution”). Rarer mutations occur later on smaller branches of the evolutionary tree. Using a DNA sequencing method, duplex sequencing, with unprecedented accuracy and sensitivity, we quantified very rare unique subclonal mutations in diagnostic specimens from five human colorectal cancers. Rarer subclones probe later evolutionary timepoints than previously possible. We confirm neutral evolution at later times and find many more subclonal mutations than expected. A novel theoretical method allowed us to extrapolate further forward in time to diagnosis. At diagnosis, every base in DNA is mutated in at least one cancer cell. In particular any therapy resistance mutation would be present.

## Introduction

Accumulation of somatic mutations is a characteristic of cancer. Solid tumors contain numerous selected clonal mutations^1^. Subclonal mutations—each present in only a small fraction of malignant cells—also contribute to phenotypic and morphologic heterogeneity within a tumor^2^, and potentially to therapeutic resistance^3^. The extent of subclonal mutations in cancer has been difficult to quantify, as the high error rate of next-generation sequencing precludes reliable detection of mutations present in fewer than 5% of cells^4^.

Here, we apply the highly accurate Duplex Sequencing (DS) methodology to quantify the extent of subclonal mutations, each present in less than 10% of the genomes, in colorectal cancers (CRCs) and adjacent “normal” mucosa. The accuracy of DS (<1 artifactual background mutation in 10^7^) bases^5^ is >10,000-fold greater than routine next-generation sequencing, enabling quantification of subclonal mutations at very high depth. Both strands of single DNA molecules are sequenced, and mutations are defined as those that are present in both strands of the same molecule at the same position and are complementary^6^. We conducted ultra-deep sequencing of 11 MSI negative CRCs (T) and adjacent normal (N) tissues. We assembled two gene libraries, a 10 kb library encoding replicative DNA polymerase active sites (5 T/N pairs), and a 13 kb library encoding genes frequently mutated in CRC (11 T/N pairs). No clonal mutations were detected in the conserved DNA polymerase genes, but clonal mutations were plentiful within the second library in tumors as expected based on TCGA data (TP53, 5/11; KRAS, 5/11; BRAF, 1/11; PIK3CA, 5/11; UMPS, 1/11)^7^.

## Results

The mean subclonal mutation frequency was 8.4 × 10^−7^±6 × 10^−8^ per nucleotide sequenced; n=11 in the tumors and 9.2 × 10^−7^ ± 1 × 10^−7^ in the “normal” colon. The mutational burden in adjacent normal tissue is not statistically different than the uncorrected mutational burden in the tumor (Fig. 1a), but when the tumor mutational burden is corrected for the percent of normal tissue present, estimated with digital imaging analysis (See SI Appendix), tumors exhibit a 1.9-fold increase in the mutational burden compared to normal tissue, which is statistically significant (Fig. 1b). The presence of numerous subclonal mutations in normal colon expands on the results of Martincorena et al^8,9^ who observed clonal mutations throughout normal esophageal tissue. There are several possible explanations for this phenomenon. First, it is difficult to compare the mutation frequencies in normal colonic tissue and relate them to relative mutation rates without knowing the corresponding proliferation histories. In normal colon stem cells divide unequally, one daughter replacing the parental stem cell while the other differentiates into the intestinal lumen; in tumors, each cell undergoes symmetric divisions. Further, it has been estimated that half of the mutational burden in a tumor was acquired in cells before birth of the founder due to the large number of cell divisions over many years before the founder cell is formed, which could also explain the observed results^10^. Finally, with age more and more “normal” cells may simply acquire a mutator mutation but still have an incomplete set of oncogenic driver mutations, whereas malignant cells may have a complete set^11^.

**Figure 1.**
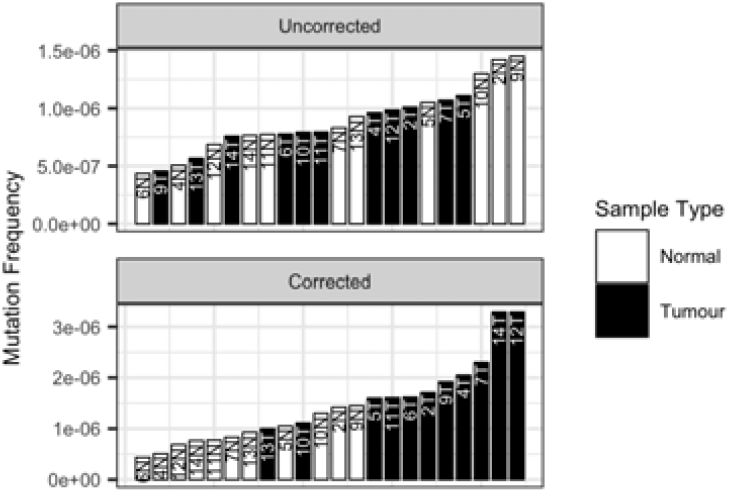
Mutation frequencies in CRCs before and after correction for the presence of normal cells. **Top. Mutation frequencies in tumor (uncorrected) and paired normal tissues obtained at a distance 1cm from the tumor margin.** Tumor and normal tissue are not significantly different by Wilcoxon rank sum test, p = 0.84. Conservative, lower tumor mutation frequencies from this figure are reported in the manuscript. **Bottom. Mutation frequencies in tumor (corrected) and paired normal tissues.** Tumor mutation frequency corrected for admixture with normal tissue. Percent normal tissue in tumor samples estimated with an automated image analysis system (SI Appendix). Tumor and normal significantly different by Wilcoxon rank sum test, p = 7.0 × 10^−5^.

We measured the mutational spectrum of the 96 possible triplets (consisting of the mutation and 3’ and 5’ bases flanking each substitution) in CRCs, adjacent normal colon, and glioblastomas (CRC data in SI Appendix, Figure S3). CRCs cluster closely in triplet space and the distribution does not differ statistically from normal colon. Cosine analysis indicates that the landscape of triplets^12^ is different in the highly conserved polymerase sequences in glioblastomas than in CRCs (SI Appendix, text, Table S6 and Figure S4) in accord with Hoang et al^13^ who found mutational spectra similar between T and N but different between different tumor types.

The contribution of selection in tumor evolution is still debated. It is generally agreed that oncogenic driver mutations conferring critical cancer phenotypes are clonal and positively selected. Some models assume successive purifying selective sweeps associated with acquisition of additional drivers^14^, while others^11,15^ assume neutral evolution after malignant transformation. The latter models featured an early mutator mutation^11,15–16^ increasing the mutation rate and accelerating the acquisition of driver mutations. In a subsequent “Big Bang model”^17^, selective sweeps also need not be invoked. Experimental studies at varying depths supported neutral evolution^13,17,18^.

Because of the branching nature of tumor evolution, increasingly rare private mutations are found in later, more numerous branches. Thus our ability to sequence more deeply with DS gives us a window into evolution further forward in time from the birth of the founder cell. In order to determine if neutral evolution continues at a constant high mutation rate at later timepoints, we sequenced multiple independent samples from five tumors to progressively increasing depths up to 20,000, using capture probes for replicative DNA polymerases. Our mathematical analysis is similar but not identical to the approach Williams et al^19^ devised to analyze the TCGA database. Both methods support the neutral model, in which most mutations do not affect fitness, and a linear relationship is predicted between the number of unique subclonal mutations and the sequencing depth up to depths approaching the reciprocal of the effective mutation rate (i.e. ca. 1.5 million, see below); this is confirmed in Figure 2 and in SI Appendix Figures S1a-e. Purifying selection and changes in growth dynamics or mutation rates could each result in deviations from linearity (see SI Appendix text and Table S4) However, this was not observed (Fig.2; SI Appendix Figures S1a-e), indicating that within the limits of our sensitivity these effects do not skew the average frequency and types of single-base substitutions in the subclonal mutational landscapes. The results suggest that neutral evolution at a high mutation rate continues as far forward in time as we can see, i.e. until the tumor reaches 20,000 cells.

**Figure 2.**
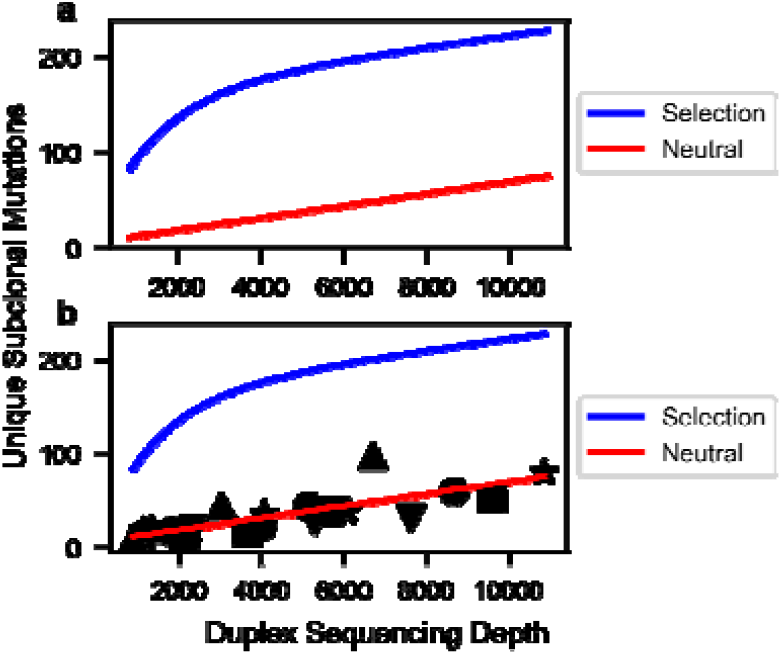
Simulated(a) and actual (b) curves for the number of nucleotide sites uniquely mutated in the DNA polymerase library (10kb) versus Duplex Sequencing depth. **a**: Simulations assume most subclonal mutations are neutral passenger mutations (solid line) or assume significant purifying selection (dashed line). Simulation methods and parameters are given in SI Appendix. Neutral model: The curve for neutral mutations alone is predicted to be approximately linear for sequencing depths up to 100,000 and after that to approach saturation in an exponential fashion. A mathematical transformation of the number of mutated sites, which is predicted to be exactly linear for all sequencing depths, is given in Methods. Selection model (blue line): This curve is the sum of curves for three classes of sites (positively selected, neutral, and negatively selected) exponentially approaching saturation of their respective sites at different rates. Positively selected sites approach saturation rapidly, while negatively selected sites approach saturation slowly, if at all. The resulting simulation shows sharp curvature. The figure is only illustrative; the curvature shown may not be detectable for weak selection (s < 0.2) combined with low mutation rate, or for a small number of positively selected sites (< 1%). **b**: Observed data super-imposed on the selected and neutral models from (a). The observed data fit the latter neutral model with correlation coefficients ranging from 0.953 to 0.999 for five tumors (shape-coded) independently sequenced at multiple depths (SI Appendix, Table S3 and Figure S1). The plotted line is the optimal regression line through all the data points from the five tumors, less precise than individual regressions for each tumor.

The slope of the lines plotted for individual tumors determine a mutation rate per template nucleotide locus per effective cell division, or “effective mutation rate” (Methods and SI Appendix) in a manner similar but not identical to Williams et al^19^. An effective cell division is defined by the addition of one new cell to the tumor, and consists of multiple actual cell divisions depending upon the differences between cell birth and death rates. Thus the “actual mutation rate” will in general be lower than the “effective mutation rate.” The observed effective mutation rate may change without a change in the actual mutation rate if the balance between birth and death rates changes during tumor growth, but our conclusions below remain robust for different growth patterns (SI Appendix). The number of effective cell divisions is based on tumor size, in that an effective cell division adds one net new cell to the tumor. The “actual mutation rate” cannot be determined without knowledge of cellular birth and death rates throughout the tumor’s history. Effective mutation rates can be used to estimate total mutational burden of a tumor (below) and to govern evolutionarily optimized therapy^20^.

Among the 5 tumors sequenced at different depths, the average effective mutation rate is 7.1 × 10^−7^. This is substantially higher than previously estimated for actual mutation rates in normal tissues based on human population genetics and/or tissue culture studies (ca. 10^−10^)^21^. However, one cannot compare an effective mutation rate to an actual one without knowledge of the proliferation history, and thus cannot definitively state whether this is indicative of a mutator phenotype.

Given a genome length of 3.1 × 10^9^, multiplying by the effective mutation rate per nucleotide, we estimate 2200 new mutations per new daughter cell added to the tumor, or for 32 doublings between the founder cell and a recently born cell when the tumor is large enough to be detected radiologically, a genetic difference of ca 65,000 subclonal mutations. In interpreting this very high number we note that the vast majority of these mutations would be private mutations detectable only by single cell or single molecule sequencing, including mutations in non-coding DNA.

## Discussion

Quantification of intratumoral genetic diversity and its relationship to therapy resistance began with the landmark 1965 use of combination therapy to prevent emergence of resistance in childhood leukemia^22^. Numerous authors have concluded that each tumor cell is genetically distinct^18,19,23–25^. Both Loeb et al^26^ and Sottoriva et al^17^ pointed out that a high mutation rate facilitates drug resistance, and Sottoriva et al^17^ made a similar point for neutral evolution. Pre-existing mutations have been linked to resistance in preclinical experimental models^27^, in single cell analysis of subclones expanded in 3D culture^25^, and in clinical cases. Thus pre-existing resistance to single agent therapy is *likely* in *many* cases.

We estimate the total burden of unique subclones in a tumor from an analysis of the unique subclones detectable at different depths by extrapolation of our data to single cell depth in tumors of different sizes (Methods and SI Appendix,). At very high depth (greater than the reciprocal of the effective mutation rate, ca. 1.5 million), the Williams et al.^19^ model and related stochastic analyses_23,28_ substantially underestimate the tumor diversity (Figures 3-5, Methods and SI Appendix). These approaches utilize the “infinite sites assumption”^29^ that any mutation is unique when first formed. This assumption requires that the number of cells in the dividing population is less than the reciprocal of the effective mutation rate (i.e., 1 million cells dividing with an effective mutation rate of 10^−10^). In these cases, there will be no mutations at the majority of sites in the population, and more than one cell with a mutation in the same site is unlikely. We have however inferred a higher effective mutation rate than previous authors with the exception of Williams et al^19^. What will be the total tumor diversity if this high mutation rate continues until the tumor reaches its minimum radiologically diagnosable size of 10^9^ cells or even beyond to the terminal phase of the disease with many more cells present in the total cancer? The infinite sites assumption is no longer applicable due to the large number of individual cell divisions per cell generation, and at each nucleotide locus we expect multiple cells will acquire the same mutation simultaneously (expected number of new cells with the given mutation = mutation rate per nucleotide locus per cell X number of cells dividing >> 1).. A mathematical approach independent of the infinite sites assumption, suitable for accurate estimation of tumor mutation burden, is given in the Methods and SI Appendix, and the predictions are confirmed in each of the five tumors sequenced at multiple differing depths (Figure 2; SI Appendix Figures S1a-e). We note that stochastic models^23,28^ and deterministic models with^19^ and without (this manuscript) the infinite site assumption, despite their different mathematical forms, give similar results over a wide range of typical experimental conditions (Figures 3 and 4; see mathematical proof in SI Appendix).. Stochastic models are more accurate than others at low cell number, and models without the infinite sites assumption are more accurate for cancers large enough to be clinically diagnosed (Figures 3 and 4, Methods, SI Appendix). The difference in predicted diversity is highly significant at diagnosis and increases dramatically as the total cancer burden increases (Figures 4 and 5).

**Figure 3:**
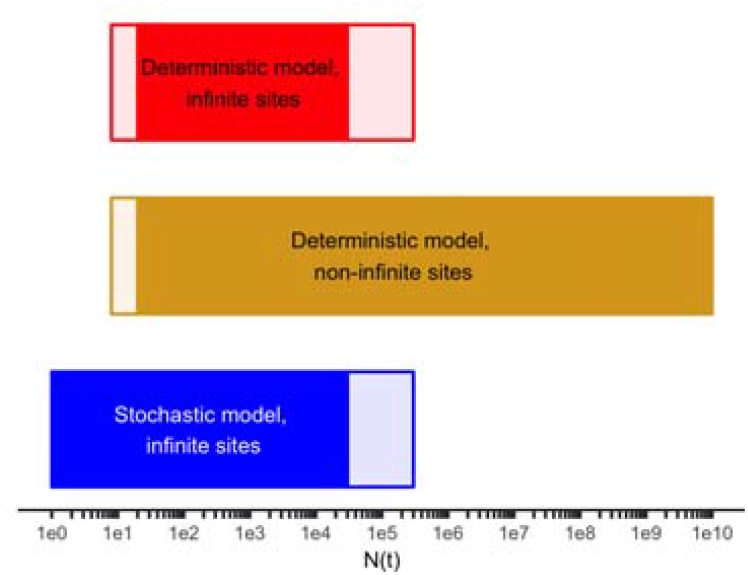
Range of applicability of various models of intratumoral diversity, vs. the number of cells in the tumor N(t) when the mutation was acquired. Sequencing to a depth D queries on average mutational events occurring when N(t) =D. Stochastic models such as Bozic et al^23,28^ are more accurate than deterministic models at early times when the tumor is small. Models without the infinite sites assumption (this manuscript) are more accurate than those with it^19,23,28^ for larger tumor masses in which the number of cell approaches or exceeds the reciprocal of the effective mutation rate. Light shaded parts of the bar represent zones where a method is not the best method but is within 10% of the best method. Solid bars indicate the method is the best method or within 1% of it. In the range of N(t) ≈ D corresponding to typical current experimental depths, all three methods are within less than 1% of each other. Parameters are b = 0.25/day, d = 0.18/day, k_mut-eff_ = 6.1 × 10^−7^.

**Figure 4:**
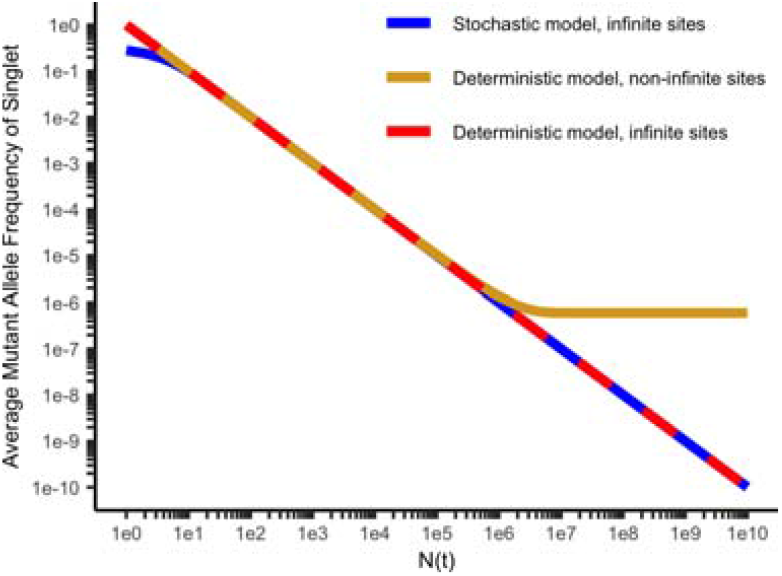
Average mutant allele frequency (MAF) for a given mutation vs N(t), the number of cells at the time it is formed, for the stochastic model with the infinite sites assumption^23,28^ (blue), the deterministic model with the infinite sites assumption^19^ (red), and without the infinite site assumption (gold; this manuscript). The deterministic model with the infinite sites assumption leads to a reciprocal relationship between N(t) and the average MAF and a straight line with a slope of −1 on a log-log plot. The deterministic model without infinite sites has an asymptotic limit for the MAF of kmut-eff for N(t) comparable to 1/k_mut-eff_ and larger. Parameters are b = 0.25/day, d = 0.18/day, k_mut-eff_ = 6.1 × 10^−7^.

**Figure 5:**
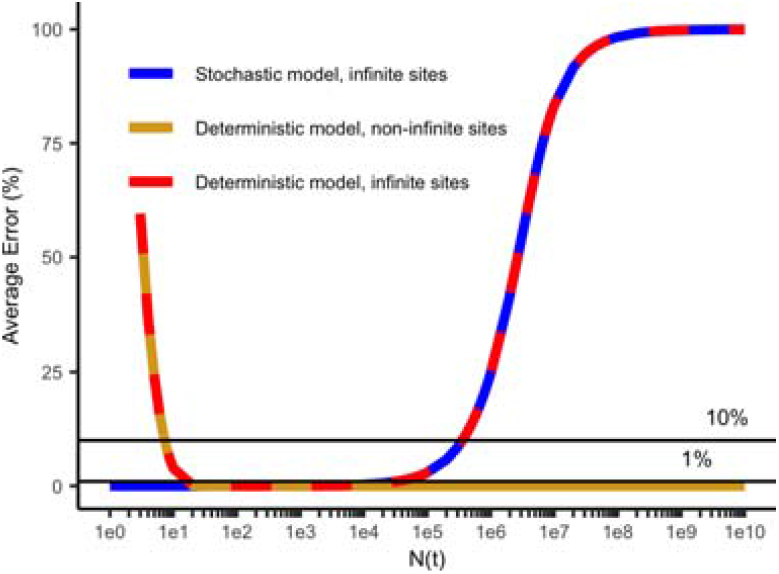
Average percentage error in determining mutant allele frequency (MAF) for each intratumoral diversity model compared to the most accurate model as a reference for the stochastic model with the infinite sites assumption^23,28^ (blue), and the deterministic model with^19^ (red) and without (gold, this manuscript) the infinite sites assumption^29^). Because the x axis is on a log scale, the errors made by models with the infinite sites assumption in estimating total tumor diversity in the high N(t) range on the right of the graph is highly significant.

In essence, the cancer enters a new biological phase when it grows to a total number of cells beyond the reciprocal of the effective mutation rate, acquiring substantially greater diversity. Under neutral evolution, the result has limited or no dependence on whether the cells are in one or numerous lesions. Subject to the assumptions that neutral evolution and a high mutation rate continue, we conclude from this analysis that every nucleotide locus is mutated in one or more cells in a clinically detectable tumor. The chance that every cell is wild type at any given neutral locus is 10^−308^. This conclusion differs from the idea that all tumor cells are genetically distinct (which could result from variation at a limited number of sites) or that pre-existing resistance is frequent or likely. Our work suggests that pre-existing resistance *to single agent therapy* is universal and inevitable.

Further, we are able to calculate the likelihood of simultaneous resistance at diagnosis to multiple *non-cross resistant* therapies pre-existing in a single cell based on mutational resistance alone, and how this probability increases as the tumor grows (Table 1, SI Appendix text and Table S5). The trend towards greater likelihood of simultaneous resistance to combinations as the tumor grows is in accord with Bozic et al 2013^28^. However, our estimates benefit from deeper sequencing at higher accuracy as well as independence from the infinite sites assumption. Optimal therapy must not only consider pre-existing resistance but also the risk of new multiply resistant subclones emerging during tumor growth^20^.

**Table 1.** Probability of emergence of multiply resistant cells. Probability P_NSCR_ that no cell in a cancer of N total cells will be mutationally resistant to all of K noncross resistant therapies, where in each case there are R neutral single bases in the genome, mutation of which confers resistance in selected scenarios. See Table S5 for additional scenarios.

| # of cells<br>(N) | # resistant<br>loci (R) | Therapies<br>(K) | $P_{\text{NSCR}}$ |
| --- | --- | --- | --- |
| $10^9$ | 1 | 2 | 0.999 |
| $10^9$ | 100 | 2 | $6.5 \times 10^{-3}$ |
| $10^9$ | 100 | 3 | $> 0.999$ |
| $10^{11}$ | 100 | 3 | 0.965 |
| $10^{12}$ | 1 | 2 | 0.604 |
| $10^{12}$ | 100 | 3 | 0.699 |

Our work has limitations (see SI Appendix) for additional discussion). It assumes that the mutation frequency in the purified gene fragments is representative of the whole genome. Mutational hotspots were not evident within our capture set.

Our data do not rule out weak selection with a selection coefficient less than 0.2 (i.e. a 20% net growth advantage per cell generation), nor can it rule out the presence of a very small number of selected sites (< 1%) (SI Appendix). This is in accord with sensitivity limits reported by Williams et al^19^.

We further assume that the effective mutation rate is a population-weighted average of different subclones and the population weighting is stable, an expected consequence of neutral evolution.

Our theoretical treatment does not consider spatial effects. However, our experimental data uses pooled DNA from 5 widely separated locations. Sottoriva^17^ concluded from multiple measurements that carcinomas are well mixed. Mixing may not affect growth and diversity under neutral evolution.

Drug resistance calculations assume that neutral evolution and a high mutation rate continue throughout a cancer’s lifetime. A continued high mutation rate is supported by high effective mutation frequencies in cell lines derived from mature cancers, and by experiments showing mutators are stably selected in mixed bacterial populations^30^. Genetic studies find that only a small minority of mutations are selected, consistent with neutral evolution throughout (SI Appendix).

Drug resistance calculations are also limited to point mutations and do not include other genetic mechanisms and non-genetic resistance mechanisms, nor do they consider that hyperploidy may provide additional sites for mutation. These additional mechanisms further strengthen our conclusion about the inevitability of pre-existing resistance to single agent therapy, while potentially increasing the number of non-cross resistant agents necessary to reliably eliminate a cancer compared to our estimate.

In summary, we have utilized DS to explore rare subclonal mutations at very high depth and accuracy in fresh frozen MSI negative CRC and neighboring normal tissue, allowing us to see further in time from tumor initiation than previously and revealing profound subclonal diversity. Our experimental data confirms a high mutation rate and neutral evolution as far forward in time as we can see. Using a novel theoretical method not confounded by the infinite sites assumption, we calculate the total tumor mutational burden assuming the high mutation rate and neutral evolution continue throughout tumor progression. We conclude, subject to this assumption, that pre-existing resistance in at least one cell to any single therapeutic agent is inevitable in any tumor large enough to be detected radiologically.

## Methods

### Specimens and DNA isolation

All participating patients gave informed consent, and all Ethical Guidelines for tissue acquisition from human participants were followed. Guidelines for study procedures were provided by the University of Utah and the Cleveland Clinic. Following surgery, tumor tissue was snap-frozen and stored at −80 °C. Complete clinical, surgical and pathological data were available for all cases. Pathology was classified according to the World Health Organisation classification (http://www.uicc.org/resources/tnm/) of colorectal tumors. All tumors were grade II. Genomic DNA was extracted using the Qiagen DNeasy Blood & Tissue Kit #69504. Histologic annotation to confirm diagnosis and cellularity of the extracted tumor was performed by frozen section microscopy of immediately adjacent tissue located within 5um. Tumor cellularity was estimated microscopically on the frozen section slides, yielding a mean adenocarcinoma cell % of greater than 50%. Microsatellite instability status was determined with BAT26 size analysis and immunohistochemical staining of MLH1, MSH2, MSH6 and PMS2^31^.

#### Duplex sequencing

Sequencing library preparation was carried out as previously described with minor modifications^6,32,33^,. See SI Appendix for further details, including capture sets, adaptor sequences, capture probes, data processing, and code availability.

### Subsampling of tumor libraries

A tumor library may be computationally subsampled in which all the DNA molecules in the library are randomly divided into sub-libraries of desired sizes, never reusing the same DNA molecule (sampling without replacement). A single library may be used to generate points along the linear curves here rather than performing separate experiments. Although we have performed separate experiments in this study, we have also shown for the future that subsampling gives valid results. See SI Appendix and SI Figures S1a-e for details.

### Evaluation of signatures

We created signatures for polDE-sequenced tumors, associated normal samples, and 5 GBM samples by the number of mutations of different types (C>A, C>G, C>T, T>A, T>G, T>C) at each different trinucleotide context (NCN or NTN). Detailed analytic methods are presented in SI Appendix.

### Determination of Mutation Rate and Mutation Burden via Sequencing the Same Sample at Different Duplex Depths

As it is not possible to sequence every genome present in a tumor, rare mutations (i.e., mutations present in one cell or a small number of cells) are infrequently sampled. Due to the branching nature of evolution, the earliest mutational events near the trunk of the evolutionary tree are scored in the majority of cells, and can be detected at low duplex sequencing depth. As we increase duplex sequencing depth, additional recent mutations that are present in a smaller fraction of the cells are also detected. As the number of nucleotides sequenced at a given genomic position is nearly always less than the number of cells in the tumor, we are unlikely to detect evidence of recent mutational events late in the tumor’s evolution. Thus, the full mutation burden in the tumor cannot be directly determined, and the estimate of mutation rates is based on an incomplete dataset.

We developed a method to estimate the mutation rate and the full mutation burden in the tumor by comparing measurements at several different duplex sequencing depths. Herein and in SI Appendix, we present the theoretical analysis for mutation rate estimation, then evaluate 5 colorectal tumors by Duplex Sequencing at a depth of up to 20,000X and an accuracy of <10^−7^.

We define the following terms and concepts (Table S2, SI Appendix):

- **Number of effective cell divisions: N_E_** the net number of times a single cell in the tumor produces a new living daughter cell to become two living cells, both of which survive. This is defined only for growing tissues like tumors. In non-malignant tissues, the total number of cells remains constant due to a balance of cell birth and death rates (b and d respectively). In a tumor b > d, and therefore the tumor grows over time. However, as d *≠* 0, it takes more than one actual cell division to make an effective cell division. That is, following any actual cell division, the daughter or parent cell may die. In such a case, that cell division does not count as an effective cell division, which refers only to cell divisions that result in an increase in cell number by one. This simple concept is convenient for analysis because, for any tumor of N cells which grew from a single founder cell, the number of effective cell divisions is, by definition, always N – 1, whereas the actual number of cell divisions is greater than or equal to the number of effective cell divisions, and cannot be determined due to the unknown history of the tumor. It is also important to distinguish cell divisions, which we have equated to the net number of all cells that have divided, from cell generations or doublings, each of which involves a very large number of individual cell divisions, the more so later in the tumor’s history.
- **Relationship between number of actual cell divisions (N_A_) and effective cell divisions (N_E_):** The rate of increase in a population due to a single cell (not including further proliferation from its daughters) is simply **b** in the absence of cell death, whereas the net rate of increase in a population in the presence of cell death is **b – d**. The ratio of these two quantities determines the relationship between actual and effective cell divisions:

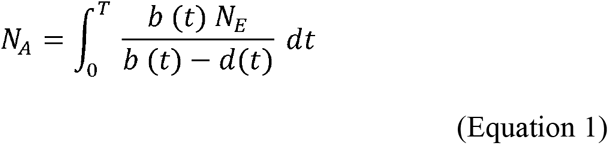 We have expressed this as a time integral since b and d may vary over time. This relationship does not apply for adult tissues without net growth, and would be mathematically undefined when **b** = **d**. Average values of **b** and **d** have been estimated using histologic techniques as the ki-67 index and the caspase 3 index, respectively. These rates are subject to both spatial and temporal heterogeneity that characterize tumor evolution, and, therefore, precise inference of the actual number of cell divisions has been difficult to obtain. Estimates of **b** for colorectal cancer have also been provided using labelled nucleotides, leading to an overall estimate of 0.25/day on average^34^. Comparing this estimate to the actual rate of increase of resistant subclones in colorectal cancer by non-invasive monitoring, an estimate for d of 0.18/day on average is obtained^3^. For example, using equation 1, if **b** and **d** have the above values, the number of actual cell divisions exceeds the number of effective cell divisions by a factor of 3.57. **Note:** the number of actual cell divisions is not utilized in the model described in this paper.
- **Duplex sequencing depth:** defined as number of duplex molecules sequenced at each nucleotide position, symbolized by **D**.
- **Number of cells in the tumor at the time of sampling: N**
- **Number of cells in the tumor at an earlier timepoint t: n(t) < N**
- **Mutation rate per nucleotide per effective cell division:** k_mut-eff_
- **Mutation rate per nucleotide per actual cell division: k_mut-actual._** Note: this parameter is not utilized in the model because it is not directly measurable.
- **Fraction of apparently unmutated single base loci at each position: F_apparent-unmutated_.** The fraction of single base loci sequenced for which there is no unique subclonal mutation detected when sequencing a tumor of N cells at depth D.
- **Reference sequence for defining unique subclones:** The consensus clonal sequence of the tumor is the reference against which unique subclonal mutations are defined. Compared to the host germline, the founder cell(s) harbors: (1) clonal mutations that were selected during carcinogenesis, and (2) random drift from the germline which occurred during the lifespan of the individual, during which the entire colonic epithelium was repopulated on a weekly basis, leading to a variation among normal colonic cells^10^. However, this prior history is not the subject of our analysis in that we only score subclonal mutations in colorectal tumors that differ from the reference sequence of the founder cell(s). Our analysis is independent of the history of the tumor prior to the birth of the founder cell.

The goal of the analysis is to infer the mutational diversity of a tumor from sequencing at a variety of depths, and to discuss the consequences for therapy. The analysis does not use the traditional input and output parameters of actual number of cell divisions N_A_ and mutation rate per base per actual cell division kmut-actual, which can only be inferred based on assumptions that are highly dependent on the unknown tumor history. Rather, it uses the input parameter of the number of effective cell divisions N_E_, which can be directly obtained from the tumor size and the corresponding output parameter of mutation rate per effective cell division k_mut-eff._ Bozic and Nowak have previously used tumor size as an approximation for the number of cell divisions^35^, as we have in this work. The other input parameters, Duplex Sequencing depth D, and fraction of apparent unmutated single base loci (relative to a tumor consensus reference) are also experimentally obtained.

We can use this analysis to characterize the mutational diversity of a tumor, including the likelihood of the presence of a mutation at an arbitrarily selected site in one or more cells of a tumor, and discuss the clinical consequences. A mathematical model for optimizing targeted therapy while accounting for tumor evolution, as parameterized by the net birth rate (birth rate minus death rate) and the effective mutation rate has previously been published. Simulations using this model demonstrate the utility of this approach, which is similar to the approach described herein^20^. However, the model cannot provide a definitive estimate of the tumor’s mutation rate per actual cell division kmut-actual, nor can it compare such a value with a comparable parameter from normal tissue to evaluate the validity of the mutator hypothesis (which states that tumors have a greater mutation rate than normal tissue). Resolution of this question requires determination of actual mutation rates, which in turn requires knowledge of the mitotic history of both the tumor and normal tissue. An illustration of the dependence of the inferred actual mutation frequency on the growth pattern is provided later in the SI Appendix. Furthermore, equation 1 is not designed to be informative for homeostatic tissues, where b = d.

### Mathematical approach

In order to model the fraction of apparently unmutated single base loci, we integrate over the entire history of the tumor, determine the fraction of apparently unmutated single base loci for daughter cells born at different times, and obtain the average fraction of apparently unmutated single base loci, weighted by the number of daughter cells born at different times. For mutations detected at a given duplex sequencing depth, the fraction of apparently unmutated single base loci is constant (independent of **n(t)**), making the average simple to calculate. At early timepoints, when there are few cells, it is less likely that a mutation will arise because there are fewer cells dividing at that time. But, if a mutation arises at this early time, it will be present in a larger fraction of the cells in the final tumor, because it is closer to the trunk of the evolutionary tree, and therefore will be more likely to be detectable. These two factors (the lower likelihood of a mutation at earlier times but greater likelihood of detecting a mutation which does occur at an earlier time) exactly counterbalance each other to give a constant number of expected detectable mutations arising at any time. These considerations lead to the following equation, which was used in our primary data analysis:

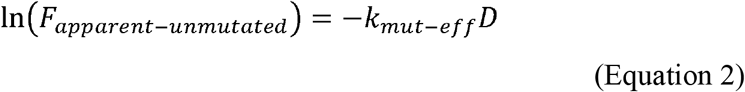

The derivation of this equation is given in SI Appendix. Plotting the natural logarithm of F_apparent-unmutated_ vs D is thus expected to lead to a straight line, with slope of -k_mut-eff_, which describes our data very accurately (SI Appendix, Figure S1a-e). The expression differs from related methods^19^ due to the absence of the infinite sites assumption. In the SI Appendix, we show that at sequencing depths below 1/k_m_ut-eff, equation 2 predicts a linear relationship between number of unique subclonal mutations and sequencing depth, similar to other methods^19^. Figure 2 is plotted in this manner due to the intuitive clarity of presentation. However, at very high depth, this simple linear relationship breaks down and greatly underestimates the total mutational burden of a tumor (Figures 3-5). These points are shown mathematically in the SI Appendix.

If we could sequence every single cell in the tumor, we could determine the total mutational burden of the tumor. In equation 2, we would substitute D = N. Raising e to the power of each side of equation 2, with D = N, we obtain:

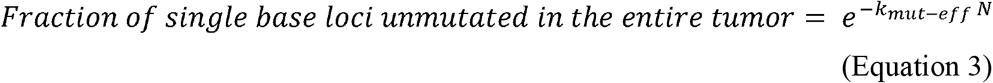

If there are R single base loci in the genome, mutation of which can lead to drug resistance to a single drug, and there are K *non-cross resistant* drugs administered, we modify equation 3 to predict the probability of no cross resistance to all K therapies, due only to mutation:

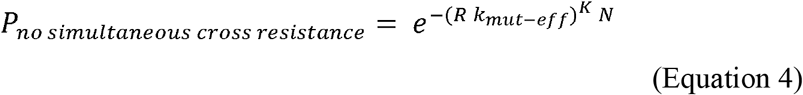

This equation was used to calculate the values in Table 1. A detailed derivation and more results of these calculations are given in SI Appendix.

In the SI Appendix, the following additional topics are considered: application of the method to the 5 individual CRC tumors sequenced at multiple depths, the relationship of this theoretical work to earlier work with particular emphasis on the mutant allele fraction as a function of sequencing depth, discussion of assumptions and approximations of the model, illustration of the dependence of actual mutation frequency on the growth pattern (eg, Gompertzian growth), evaluation of the likely consequences for the estimate of tumor mutational burden and for the mutator hypothesis, and simulations defining the sensitivity of the method for detecting deviations from neutral evolution.

## Acknowledgments

Research reported in this publication was supported by the National Cancer Institute of the National Institutes of Health under award numbers NCI P01-CA077852, and NCI R01-CA160674 (L.A.L.). The content is solely the responsibility of the authors and does not represent the official views of the National Institutes of Health. Initial studies on mutation frequency as a function of sequence depth were carried out by Edward Fox. We thank: Michael Schmitt, Jared Roach, James Shen, and for comments and Tom Walsh and Ming Lee for assistance with sequencing. De-identified samples were collected under an IRB on file with the University of Utah, in compliance with U.S. Code of Federal Regulations, 45 CFR Part 46. Raw sequencing data was deposited in the SRA under accession number SRP135906.

