## Supplemental Information for "Unexpectedly High Subclonal Mutational Diversity in Human Colorectal Cancer and Its Significance"

### Supplementary Methods

#### Capture sets.

We used two different capture sets in this analysis; first, a capture set consisting of CRC-specific targets (capture set of genes mutated in CRC); second, a capture set consisting of Polδ1 and Polε (capture set of polymerase genes, pol DE capture set). Probes for these capture sets were ordered from IDT, and sequences are provided in Dataset S1.

#### Duplex sequencing library preparation and capture.

Sequencing library preparation was carried out as previously described with minor modifications[1,2]. Adaptor sequences were prepared as described therein. In brief, DNA was sonicated, end-repaired, and A-tailed, and ligated to duplex-sequencing adapters using a 10× molar excess of adapters (MWS51 and MWS55; Table S1). Following binding and elution cleanup with 0.8 volumes of AMPure XP beads (Agencourt), the adaptor-ligated DNA was PCR-amplified for 9 cycles with the KAPA Biosystems hot-start high-fidelity kit, using primers MWS13 and MWS20. Following purification with AMPure XP beads, iterative capture was performed as described[333]. Between 250 and 1000 ng of genomic DNA was used per sequencing library, with up to 7 μg of DNA per sample to achieve varying sequencing depths. Sequences of capture probes used (Integrated DNA Technologies) are given in Table S2. Final PCR-amplification was performed using primers MWS13 and MWS21 for 5 or 6 cycles. Sequencing was performed on a HiSeq2500.
The major limitation of duplex sequencing (and ultradeep sequencing, in general) is the cost of repeatedly sequencing thousands of reads. As a result, focused portions of the genome are enriched using iterative capture[3] rather than interrogating the entire genome.

#### Subsampling of tumor libraries.

A tumor library may be computationally subsampled in which all the DNA molecules in the library are randomly divided into sub-libraries of desired sizes, never reusing the same DNA molecule (sampling without replacement). A single library may be used to generate points along the linear curves here rather than performing separate experiments. In figures S1a-e , we display the regression line from the experimental libraries (black points line, and shaded 95% confidence interval determined by a 2-sided t test with N-2 degrees of freedom, where N is given in Table S3), as well as a set of libraries generated by computationally merging the experimental libraries and then computationally splitting them into random sub-libraries of similar sizes to the original experimental libraries (red points and symbols). We see that the random sub-libraries from a merged library generate equivalent curves to separate experiments, for each tumor. We show only one instantiation of random subsampling, but this procedure was performed four times for each tumor with similar results. In 19/20 cases (95%) the subsampled regression line is within the 95% confidence interval of the regression line determined by separate experiments. The exception is shown in Figure S1e.

#### Evaluation of signatures.

We created signatures for polDE-sequenced tumors, associated normal samples, and 5 GBM samples by the number of mutations of different types (C>A, C>G, C>T, T>A, T>G, T>C) at each different trinucleotide context (NCN or NTN). These signatures were normalized by dividing the count of mutations of a given type at a given context by the number of times that context was sequenced. Mutation signatures were compared using cosine similarity, as described in Alexandrov et al.[4], and comparisons were compared using the aov function in R, with posthoc analysis performed using the TukeyHSD function. Results of these analyses can be found in Figures S3 and S4, and in Table S6.

#### Code availability.

Software for Duplex Sequencing is available at https://github.com/loeblab/Duplex-Sequencing/tree/Duplex-V-1.2.

**Data processing.**
Processing of Duplex Sequencing data was performed essentially as previously described. Data from this paper has been uploaded to the Sequence Read Archive under accession SRP135906. Subclonal mutations are defined as mutations present in fewer than 10% of sequencing reads. Mutations are scored if complementary and present in both sense and antisense strands at the same position in individual DNA molecules. The frequency of subclonal mutations represents the total number of subclonal mutations detected divided by the total number of nucleotides assayed at all positions. To obtain multiple depths for each sample, we merged different libraries, with each library being used once. PicardTools CollectHsMetrics was used to determine mean depth, and the CountMuts.py program published with the DS pipeline to count numbers of unique positions mutated.

**Determination of Mutation Rate and Mutation Burden via Sequencing the Same Sample at Different Duplex Depths.**
As it is not possible to sequence every genome present in a tumor, rare mutations (i.e., mutations present in one cell or a small number of cells) are infrequently sampled. Due to the branching nature of evolution, the earliest mutational events near the trunk of the evolutionary tree are scored in the majority of cells, and can be detected at low duplex sequencing depth. As we increase duplex sequencing depth, additional recent mutations that are present in a smaller fraction of the cells are also detected. As the number of nucleotides sequenced at a given genomic position is nearly always less than the number of cells in the tumor, we are unlikely to detect evidence of recent mutational events late in the tumor's evolution. Thus, the full mutation burden in the tumor cannot be directly determined, and the estimate of mutation rates is based on an incomplete dataset.

We developed a method to estimate the mutation rate and the full mutation burden in the tumor by comparing measurements at several different duplex sequencing depths. Herein, we present the theoretical analysis for mutation rate estimation, then evaluate 5 colorectal tumors by Duplex Sequencing at a depth of up to 20,000X and an accuracy of $<10^{-7}$. We then present additional methods for evaluating the full mutation burden of a tumor, and theoretical implications for mutational drug resistance. Finally, we evaluate the sensitivity of our methods for distinguishing neutral evolution from other mechanisms.

Please see Methods and Table S2 for a definition of terms and concepts.

### Explanation and Discussion of Modeling

In the subsequent sections, the following topics are considered: (1) presentation of the mathematical approach, (2) application of the approach to human colon cancers sequenced at multiple depths, (3) Methods for estimating mutation frequencies for tumor masses large enough to be clinically diagnosable (4) comparison to the related mathematical approaches of Williams et al[5] and of Bozic et al[6,7] (5) discussion of assumptions and approximations of the model , (6) illustration of the dependence of actual mutation frequency on the growth pattern, and evaluation of the likely consequences for the estimate of tumor mutational burden and for the mutator hypothesis, (7) discussion of the likelihood of existing resistance to one or multiple non-cross resistant agents in malignant cells during tumor growth, and simulations of the sensitivity of this method for detecting weak selection.

**Mathematical approach.**
In order to model the fraction of apparently unmutated single base loci, we integrate over the entire history of the tumor, determine the fraction of apparently unmutated single base loci for daughter cells

born at different times, and obtain the average fraction of apparently unmutated single base loci, weighted by the number of daughter cells born at different times. For mutations detected at a given duplex sequencing depth, the fraction of apparently unmutated single base loci is constant (independent of **n(t)**), making the average simple to calculate. At early timepoints, when there are few cells, it is less likely that a mutation will arise because there are fewer cells dividing at that time. But, if a mutation arises at this early time, it will be present in a larger fraction of the cells in the final tumor, because it is closer to the trunk of the evolutionary tree, and therefore will be more likely to be detectable. These two factors (the lower likelihood of a mutation at earlier times but greater likelihood of detecting a mutation which does occur at an earlier time) exactly counterbalance each other to give a constant number of expected detectable mutations arising at any time. Specifically, at any given time **t**, we have (# symbolizes "number"):

$$Expected\ \#\ of\ cells\ with\ new\ mutations\ at\ a\ given\ nucleotide\ for\ the\ totall\ of\ all\ cells\ born\ at\ time\ t = k_{mut-eff}\, n(t)$$

(Equation S1)

In equation (S1), we assume that the n(t) cells present at time t undergo effective and synchronous divisions, eventually creating a new cell generation of n(t) daughter cells. This may require more than n(t) actual cell divisions. Note that expectation values need not be integers, and unlike probabilities, are not restricted to values, between zero and one.

$$Fraction\ of\ malignant\ cells\ in\ the\ final\ tumor\ that\ will\ arise\ from\ a\ single\ cell\ born\ at\ time\ t = \frac{1}{n(t)}$$

(Equation S2)

In equation S2, we consider the branching nature of tumor evolution, and the fact that cells born in a larger tumor are further out on branches.

$$Expected\ \#\ of\ mutations\ detected\ from\ a\ cell\ born\ at\ time\ t = \frac{D}{n(t)}$$

(Equation S3)

Equation S3 follows from equation S2, and the fact that we are randomly sampling the sequences of cells from the tumor at depth D.

Multiplying equation S1 by equation S3, we find:

$$Expected\ \#\ of\ mutations\ detected\ at\ a\ single\ base\ arising\ at\ time\ t = k_{mut-eff}\, D$$

(Equation S4)

Further, from equation S4 and the zero term of the Poisson distribution, it follows that the fraction of apparently unmutated single base loci $F_{apparent\text{-}unmutated}$ from mutations arising in cells born at any time t is given by:

$$F_{apparent-unmutated}\ from\ cells\ born\ at\ time\ t = e^{-k_{mut-eff}\, D}$$

(Equation S5)

Since (S5) is independent of time, a weighted average over the history of the tumor is given by the same expression. When we take the population-weighted average of equation S5, we integrate over all times in the tumor's history. Since every cell in the tumor was born at some point in the tumor's history, this integral over time is equivalent to integrating over all cells in the tumor. The average is then the time integral of (S5) over 0 to T, divided by T:

$$Average\ F_{apparent-unmutated} = e^{-k_{mut-eff} D}$$

(Equation S6)

**Method for estimating $k_{mut-eff}$ from sequencing at multiple duplex depths.**
Taking the natural logarithm of both sides of (S6), we obtain:

$$\ln\left(F_{apparent-unmutated}\right) = -k_{mut-eff}\ D$$

(Equation S7)

Thus, plotting the natural logarithm of the fraction of apparently mutated single base loci for several different duplex sequencing depths should yield a straight line with slope – $k_{mut-eff}$. At duplex depth (D) = 0, no mutations will be observed and the fraction of apparently unmutated single base loci will be 1, leading to a natural logarithm of 0. Thus, the origin (0, 0) should also be on the line.

**Fraction of unmutated single base loci in every cell in the tumor.**
If we could sequence every single cell in the tumor, we would determine the actual value of this quantity. Thus, this number is given by equation S6, when D = N:

$$Fraction\ of\ single\ base\ loci\ unmutated\ in\ the\ entire\ tumor = e^{-k_{mut-eff} N}$$

(Equation S8)

**Fraction of single base loci which are newly mutated in m cells in the tumor.**
From equation S8 and the Poisson distribution, we can define this quantity:

*Fraction of single base loci newly mutated in exactly m cells in the tumor*:

$$\frac{(k_{mut-eff} N)^m\ e^{-k_{mut-eff} N}}{m!}$$

(Equation S9)

**Application to 5 tumors sequenced at multiple depths.**
The analysis above predicts that a plot of $\ln(F_{apparent-unmutated})$ vs D will be linear, with a negative slope equal to $-k_{mut-eff}$, and will go through the origin although this point was not entered into the regression (all sites apparently unmutated at a sequencing depth of zero).

We find this is indeed the case for the mismatch repair proficient colorectal cancers that were sequenced at 4 depths ranging up to 20,000 (Figures S1a-e, Table S3) ). Linear correlation coefficients $R^2$ ranged from 0.953 to 0.999, with a mean +/- 1 standard error of 0.974 +/- 0.017. The percent of variation in data explained by the linear model is given by 100 times the square of the correlation coefficient, and ranged from a low of 90.8% in one tumor to a high of 99.9%, with a mean +/- 1 standard error of 94.9 +/- 3.3 %.

Only one of 5 estimated y-intercepts is statistically different from zero according to a 2-sided 95% confidence interval using the student's t-test with 2 degrees of freedom, and, for that one, the lower boundary of its 95% confidence interval was approximately 15% of the smallest data-point value. The mean+/- one standard error of the estimated y intercepts over the 5 tumors is -4.3 +/- 5.5 X $10^{-4}$, indicating that the overall estimate of the y intercept is less than one standard deviation from zero. The excellent agreement of data with the model using five separate tumors renders more complex models less likely.

The estimated effective mutation rates $k_{mut-eff}$ for the 5 tumors ranges from 2.8 X $10^{-7}$ to 1.5 X $10^{-6}$. The mean effective mutation rate (+/- 1 standard error) over 5 tumors is 7.1 +/- 4.4 X $10^{-7}$ per base per effective cell division. For a genome of 3.1 X $10^9$ bases, this means that each surviving new cell that adds to the tumor population has approximately 2,200 new mutations compared to its parent. Since these mutations are occurring randomly throughout the genome, only 1% of which is coding, this amounts to approximately 22 new mutations in coding segments per daughter cell, some of which will be synonymous. Many of these mutations will be private and only detectable if sequencing to single cell depth.

**Relationship to earlier work[5-13].**

There have been many attempts to model tumor progression. Beckman and Loeb[10] and Beckman[11] modelled carcinogenesis with selection operating only on dominant and recessive oncogenes, with neutral evolution after the formation of the founder cell. Sottoriva et al[12] investigated in depth the similar hypothesis that colorectal cancers grow as a single expansion with most selected mutations occurring early. Multiparameter profiling of individual clones from diverse locations within these cancers demonstrated an absence of selective sweeps. Williams et al[5] analyzed the TCGA database and found remarkable subclonal diversity and evidence for neutral evolution. Martincorena et al[9] also found strong support for neutral tumor evolution in a comprehensive analysis of the ratios of synonymous to nonsynonymous mutations.

Herein, we compare and contrast the proposed models and methods.

The Williams et al.[5] modelling approach and ours are similar in a number of important respects:

- Both use quantitative analysis of rare mutations to support the theory of neutral evolution.
- Both parameterize the model in terms of observables such as effective cell divisions, rather than on parameters such as actual cell divisions, that are difficult to infer.
- Both predict a linear relationship between sequencing depth and a quantity related to observed genetic diversity, wherein the slope reveals the effective mutation rate, i.e. $k_{mut-eff}$.
- For all observations at depths of order of magnitude less than the reciprocal of the mutation rate, (in our case less than the order of 1.4 X $10^6$), the predictions of the two approaches are identical. A mathematical explanation of why this is the case is given at the end of this section.
- Both models conclude that significant subclonal diversity might be a source of pre-existing drug resistance.

Our approach and that of Williams et al.[5] approach differ in several important respects:

- Purpose: Williams et al.[5] examined the TCGA database at low depth utilizing a variety of techniques with varying accuracy, and described the statistical distribution of mutation frequencies in whole exome sequencing within tumor types. The TCGA database scores primarily for clonal mutations. In contrast, we isolated DNA from surgical specimens and used an assay that is 10,000-fold more accurate than standard NGS to examine rare subclonal mutations at depths of up to 20,000X.
- Representation of distribution of variants as a function of their frequency: Williams et al.[5] implicitly assume that any new mutation occurring at time *t* is occurring in **only one copy, rather than being born new in multiple copies in different cells**: "for a new mutation occurring at any time *t*, its allelic frequency (relative fraction) *f* must be the inverse of the number of alleles in the population." This quote drives equation (5) in the Williams et al.[5] paper and the rest of the analysis. This assumption, that any new mutation at a particular site occurs uniquely, rather than simultaneously in multiple cells, is termed the "infinite sites approximation."[13] The assertion is valid for depths less than one over the mutation rate, i.e. the

subject of the Williams et al.[5] analysis and our experimental work. In contrast, our equation (S1) describes the expected number of new mutations simultaneously within different cells as the product of a mutation rate and the number of cells dividing simultaneously. **This is an average and does not have to be an integer like 0 or 1.** The probability that there will be any given integral number of DNA molecules with the variant of interest is then given by the Poisson distribution. Below, we will derive the predictions of the mutant allele frequency as a function of the number of alleles in the population for our model, **which does not rely on the infinite sites approximation,** and contrast it with other models.

- Difference between the methods' predictions at depths less than one over the mutation rate, i.e. variant frequencies greater than the mutation rate: in this range, the two methods give very similar predictions. This is because at these depths, it is unlikely that there will be more than one copy of a given new mutation being formed at any instant. When $k_{mut\text{-}eff}\, n(t) << 1$, most sites in the genome will remain unmutated, and a small minority of sites will have only one copy of a new mutation formed at that instant. A very small minority of sites will have 2 or more copies of the new mutation simultaneously forming, and thus the Williams et al.[5] approximation is accurate in this range (Figure 3).
- Difference between the methods' predictions at depths much larger than one over the mutation rate: in this range, the Williams et al.[5] approach, which asserts that the variant frequency will be inversely proportional to the depth at all depths, differs from ours. Given the mutation rate of approximately $7 \times 10^{-7}$ per effective cell division, and the fact that the smallest tumor visible on CT will contain between $10^8$ and $10^9$ cells[14], the product $k_{mut\text{-}eff}$ times n(t) will be >> 1 before the tumor is diagnosed and this will only increase as the tumor grows. In such cases, we do not believe that the variant frequency will be 1/n(t) (or 1/D, given that at depth D we are on average observing mutations that occurred when n(t) =D) as asserted by Williams et al.[5]. Rather the expected number of copies of a new mutation formed at a doubling from n(t) to 2n(t) will be as given in equation (S1), i.e. $k_{mut\text{-}eff}\, n(t)$. For the case of the tumor with $10^9$ cells, if they all divide approximately simultaneously, we expect approximately $(7 \times 10^{-7}) \times 10^9$ or 700 of the newly formed $10^9$ daughter cells to contain the same new variant created approximately simultaneously. The chance of only one copy of the new variant being created under these circumstances is negligible. Thus, at high n(t), the variant frequency does not continue to decrease as 1/n(t), but is given by:

$$variant\ frequency\ for\ cells\ born\ at\ time\ t = \frac{expected\ number\ of\ copies\ of\ variant\ formed}{number\ of\ cells\ dividing\ at\ time\ t} = \frac{k_{mut-eff}\, n(t)}{n(t)} = k_{mut-eff}$$

(Equation S10)

Thus, the curve of variant frequency versus depth predicted by the two methods is very different in this region (Figure 4). The Williams et al.[5] curve continually decreases with increasing depth with an asymptotic limit of zero, whereas the model we propose has an asymptotic limit of $k_{mut\text{-}eff}$.

In the region where $k_{mut\text{-}eff}\, n(t) \approx 1$, there is a transitional zone and the predictions begin to diverge. To describe the entire curve precisely, we utilize *conditional* expectation values for the expected number of mutant alleles simultaneously arising at particular site at a time t, when n(t) = D cells are each undergoing individual divisions, rather than utilizing an infinite sites approximation. (Note throughout our treatment and that of Williams et al.[5], the assumption is that at a depth D we are *on average* sampling cells formed at time t where n(t) =D). The term *conditional* means we will apply the condition that at least one mutant allele has been experimentally observed at the site in question. At very low depth relative to the reciprocal of the effective mutation frequency, most sites will have no mutant alleles, a small minority will have one mutant allele, and increasingly smaller minorities will have higher integral numbers of mutant alleles, leading to a non-integral average as described above. At these low depths, if a single mutant allele is observed the number of mutant alleles cannot be zero, and one is a much more likely value than higher integral values. Thus, the infinite sites approximation, which assumes that the mutant allele number is in fact one wherever mutations are observed, is effective in this range. If the mutant allele number is one, the mutant allele fraction will be 1/n(t) as asserted by Bozic et al.[6,7] and Williams et al.[5].

To develop an expression that will be accurate at all depths, we start with an exact expression for the total number of mutant alleles expected to be newly formed at a particular site at time t, which is a probability weighted average of all possible integral values from 0 to n(t), where $p_n$ is the probability that there are n simultaneously formed copies of the mutant allele:

$$Expected\ number\ of\ new\ mutations\ at\ a\ particular\ site\ at\ time\ t = \frac{\sum_{n=0}^{n(t)} n * p_n}{\sum_{n=0}^{n(t)} p_n}$$

(Equation S11)

The denominator is equal to 1, since it represents the sum of the probabilities of all mutually exclusive possible outcomes. Thus,

$$Expected\ number\ of\ new\ mutations\ at\ a\ particular\ site\ at\ time\ t = \sum_{n=0}^{n(t)} n * p_n$$

(Equation S12)

But comparing to equation S12 to equation S4, we see that:

$$\sum_{n=0}^{n(t)} n * p_n = k_{mut-eff} D$$

(Equation S13)

We can now compute the *conditional* expected value of the number of mutant alleles newly formed simultaneously at a particular site *given that this number is greater than or equal to one,* i.e. at sites where there has been at least one mutation. We take a weighted average again of integral values of 1 or greater, not including zero.

$$Conditional\ expected\ number\ of\ mutant\ alleles\ at\ a\ given\ site\ given\ that\ there\ is\ at\ least\ one\ mutant\ allele = \frac{\sum_{n=1}^{n(t)} n * p_n}{\sum_{n=1}^{n(t)} p_n}$$

(Equation S14)

We note that the sum in the numerator, from 1 to n(t), is the same as the sum from zero to n(t), since the zero term is zero. Further the denominator is equal to the sum from zero to n(t) (previously noted to be 1) minus the zero term, the probability of no mutant alleles, or $p_0$. Making these substitutions, we have:

$$\text{Conditional expected number of mutant alleles at a given site given that there is at least one mutant allele} = \frac{\sum_{n=0}^{n(t)} n * p_n}{1 - p_0}$$

(Equation S15)

Substituting the value of the numerator from equation S13, and the value of p0 from equation S6 ($p_0$ and average $f_{\text{apparent unmutated}}$ are synonyms), we obtain:

$$\text{Conditional expected number of mutant alleles at a given site given that there is at least one mutant allele} = \frac{k_{mut-eff}\, D}{1 - e^{-k_{mut-eff}\, D}}$$

(Equation S16)

To derive the conditional expectation of the mutant allele fraction, we divide by n (t), setting n(t) = D as before:

$$\text{Conditional expected mutant allele frequency at a given site given that there is at least one mutant allele} = \frac{k_{mut-eff}}{1 - e^{-k_{mut-eff} D}}$$

(Equation S17)

To evaluate this expression at depths much lower than the reciprocal of the effective mutation frequency, we expand the exponential in a Taylor's series. Taylor's theorem states that any continuous function f(x) may be expanded in the region around f(a):

$$f(a + \Delta a) = f(a) + \Delta a\, f'(a) + \left[\frac{(\Delta a)^2}{2!}\right] f''(a) + \left[\frac{(\Delta a)^3}{3!}\right] f'''(a) \ldots$$

(Equation S18)

where f', f'', and f''' are successive derivatives of f.

At depths much lower than the reciprocal of the effective mutation frequency, the exponent is nearly zero, and the exponential term may be approximated by a Taylor's series with $f(a) = e^a$, $a = 0$, $\Delta a = -k_{mut\text{-}eff}\, D$, truncated at the linear term, i.e.

$$e^{a+\Delta a} = 1 + \Delta a + \frac{(\Delta a)^2}{2!} + \frac{(\Delta a)^3}{3!} + \cdots \cong 1 + \Delta a, \textit{when } \Delta a \ll 1$$

(Equation S19)

Applying this approximation to equation S17, we see that at low depth the conditional expected average mutant allele frequency is equal to $1/D \approx 1/N(t)$, in agreement with Williams et al.[5].

But at high depth, where D is greater than the reciprocal of the mutation rate, the mutant allele frequency does not approach zero as in the Williams et al.[5] formulation. Rather, examining equation S17, the exponential term approaches zero, and the conditional expected average mutant allele frequency approaches $k_{mut\text{-}eff}$.

Williams et al.[5] have performed a simulation and compared it successfully to their mathematical treatment. This shows that the mathematics and simulation are mutually consistent. However, the simulation utilized only $10^6$ individual cells, the technical limit of such stochastic simulations, and within the range of validity of the infinite sites assumption. The model results were then "scaled up" using the same assumption, to a full-size tumor. The infinite sites assumption is, in our opinion, not applicable in that range, and therefore one cannot use it to evaluate clinically detectable tumors.

The underestimate of the total burden of mutations in the tumor associated with the infinite sites assumption is given by the area between the two predicted curves of mutation frequency versus depth at high sequencing depth >> $1/k_{mut\text{-}eff}$ (Figure 4). Although the difference in asymptotic limits in the y axis is small, equal to $k_{mut\text{-}eff}$, the x-axis width of this region is three orders of magnitude greater at diagnosis ($10^9$ cells) than the region of validity of the infinite sites assumption (up to $10^6$ cells). Moreover, as the malignancy grows and spreads, it could contain as many as $10^{12}$ cells in the terminal phase. Thus, the area between the curves represents a quantitatively significant difference in the predicted diversity with and without the infinite sites assumption (Figure 5).

- Consequence of the two models' divergent predictions at very high depth: a theoretical estimate of the total mutation burden within the tumor, and a calculation of the extent of drug resistance to one drug or to multiple non-cross resistant drugs due to mutation(s) within the same cell, as given in this work, requires a model which is accurate in the range $k_{mut\text{-}eff}\, n(t) >> 1$. Due to the ever-increasing number of *cell divisions* per *cell generation* as the tumor grows, the discrepancy between the two models is quantitatively significant when the problem of the total mutational burden within the tumor is considered.
- Mathematical form of the predictions: although the Williams et al.[5] formulation and ours both predict linear relationships of genetic diversity as a function of sequencing depth, the measure of genetic diversity on the y axis differs. In Williams et al.[5], the y axis is the intuitive quantity of number of unique subclones. Because this approximate treatment is accurate within the depth range of our experiments, and is intuitive, we have presented it this way in the main text. However, our more general formulation presented herein has the y axis as the natural logarithm of the unmutated fraction of bases sequenced, i.e. ln(1-x) where x is the fraction of the capture set for which a variant sequence is detected. Moreover, the presentation of Williams et al.[5] has a positive slope proportional to the effective mutation rate, whereas ours has a negative slope equal to the effective mutation rate, due to the different quantity on the y axis.
- Near equivalence of the two formulations at low depth: we have given above both a qualitative argument and a mathematical argument for the near equivalence of the formulations at low depth, the latter focusing on the mutant allele frequency curve as a function of depth. We now will again show this near-equivalence at low depth, this time focusing on the curve of observed unique subclonal mutations as a function of depth. We begin with equation S7, noting that $F_{apparent\text{-}unmutated} = 1 - x$, where we use x to denote the fraction of sites within the capture set at which at least one copy of the mutation has been observed. Thus, equation S7 becomes:

$$\ln\ (1-x) = -k_{mut-eff}\, D$$

(Equation S20)

Mathematical equivalence at low depth is shown as follows:

Setting f(x) = ln (1-x), a =1 and Δa = -x, we expand ln(1-x) as a Taylor series (Mathematical Tables from the Handbook of Chemistry and Physics, Chemical Rubber Company Publishing, Cleveland, Ohio, 1936, pp. 278-279) in powers of x about 1 , yielding:

$$\ln(1-x) = -x - \frac{x^2}{2} - \frac{x^3}{3} - \cdots$$

(Equation S21)

At low depth, x << 1, since variants will be detected only in a minority of the capture set loci. Thus, the higher order terms in x are negligible, and we have

$$\ln(1-x)\ \approx\ -x\ for\ 0 < x \ll 1$$

(Equation S22)

But x, the fraction of sites with a variant observed, is proportional to the number of observed unique variants, so that plugging equation S22 into equation S20, we see that the number of observed unique variants is nearly exactly proportional to the depth, at low depths, as asserted by Williams et al[5]. In summary, our model and that of Williams et al.[5] give indistinguishable predictions at depths less than $1/k_{mut-eff}$, but differ in important ways at higher depths representing three orders of magnitude of further tumor growth before the tumor grows to the minimal size at which it can be diagnosed clinically.

Bozic et al.[6] use a stochastic model to assess neutral evolution, and derive a correction to the Williams et al.[5] formula, which is significant at low depths (10-100) representing subclones with an apparent frequency of $10^0$ to $10^{-2}$. Bozic et al.[6] validate their approach using data in this range. Stochastic effects are more prominent with small numbers of cells, and the Bozic et al. model[6] will be more accurate in this very low depth range than either the Williams et al.[5] approach or ours. The Bozic et al[6] model gives two correction factors. The first 1/(1- δ) is equal to the b/(b-d) term which normalizes between the actual and effective mutation rates, and is present in both the current work and the Williams et al.[5] formulation. The second correction is factor of $(1-(d/b)^{n(t)+1})$ which is significant at low n(t).

Importantly, however, the Bozic[6] treatment, like Williams et al.[5], uses the infinite sites approximation and is thus not applicable when the tumor cell number increases beyond the reciprocal of the effective mutation rate, or at sequencing depths greater than this number.
A chart of the regions of validity of the stochastic infinite sites model[6] the continuous infinite sites model[5] and the model herein is provided in Figure 3.

We have compared our methods and work to earlier related work, particularly Williams et al.[5] and Bozic et al.[6]. Our independently developed work has striking similarities to but important differences from these earlier studies. Importantly, all three methods give the same predictions in the range of DNA sequencing depths currently explored experimentally, with the exception of very low depths, where the stochastic treatment of Bozic et al.[6] is more accurate. The other methods utilize the "infinite sites assumption"[13], which assumes that any mutation is unique at the instant it is formed. This assumption is true as long as the product of the effective mutation rate and the number of cells comprising a cell generation is much less than one. That condition is met for all experimental work done to date, as well as the conditions of all prior simulations. However, our experimental results document a higher effective mutation rate than previously considered, of the order of $10^{-6}$. We then consider the "thought experiment", currently not feasible, in which every cell in the smallest diagnosable tumor mass, or $10^9$ cells, is sequenced to determine the total mutation burden of such a mass. Under these conditions, the "infinite sites assumption" will be violated, since it is violated in all cell generations involving on the order of $10^6$ cells or higher. Our method is unique in that it accurately quantifies the mutation burden in this situation, because it is independent of the "infinite sites assumption" (Figures 3-5). This allows us to make more accurate and definitive statements about drug resistance than was previously possible. We conclude that within a tumor mass of $10^9$ cells, no DNA locus will be wild type in every cell unless mutations at that locus are lethal. While many authors have stated that each tumor cell is distinct (which could be achieved with diversity at a limited number of sites), and that pre-existing resistance to a single therapy is *likely to be common*, we can now say that pre-existing resistance to a single therapy in at least one cell from single base substitutions alone is *universal* and *inevitable.*

**Discussion of assumptions and approximations of the model[5,9,15-25]**

We discuss the following caveats to the proposed model: (1). The proposed model assumes that most mutations are neutral, (2) the proposed model assumes an average effective mutation frequency $k_{mut\text{-}eff}$ can be calculated across subclones, which may be mutating at different rates, (3) the proposed model estimates the total mutational burden of the tumor by extrapolation from the genes sequenced to the entire genome, and (4) the proposed model estimates the total mutational burden of the tumor by extrapolation from the maximum sequencing depth to a much larger number: i.e. the total number of cells in the tumor.

*Assumption of neutrality.*

We have observed a linear curve of the logarithm of the fraction of apparently unmutated bases as a function of sequencing depth. If selection associated with partial or complete clonal sweeps reduced or eliminated subclonal diversity, the absolute value of the slope should decrease with increased sequencing depth, resulting in upward curvature (less negative slope), contrary to observation. Moreover, this upward curvature would produce linear regressions with negative y-intercepts, again contrary to observation. Thus, there is no evidence from our experimental data for deviation from neutrality.

A variety of experimental and theoretical studies support the notion that, while certain key sites are highly selected in carcinogenesis, the majority are neutral. A recent population dynamic analysis of a large number of microscopic sectors of a hepatocellular carcinoma biopsy was found to be wholly consistent with a neutral model[15]. An analysis of mutation burden of tumors as a function of depth using the TCGA database was published which strongly supports a neutral evolutionary model for colorectal cancer[5]. Moreover, a theoretical study of driver and passenger mutations, supported by bioinformatics analysis of observed mutations, has concluded that neutral passenger mutations are more numerous[25], and neutral Poisson models have been previously employed in the analysis of divergence between metastases and primary[17], and in the timing and heterogeneity of resistance mutations[18].

A comprehensive survey of nonsynonymous/synonymous mutation ratios adjusted for tissue and site-specific mutation patterns found a maximum of 1,230 positively selected genes in cancer at the 0.05 level of significance, and when the significance threshold is adjusted to reduce the false detection rate due to a large number of statistical comparisons, the number of definitively positively selected genes reduced to 45[20]. This amounts to anywhere between 0.15% and 4% of the approximately 21,000 genes in the human genome. Even within a positively selected gene, it is likely that no more than 28% of the amino acids within the corresponding protein affect function when mutated[21]. Overall, therefore, somewhere between 0.04% and 1% of genomic bases are expected to be positively selected in cancer.

Regarding negative selection, Zhou et al. found between 16 and 326 genes affected, depending on the significance threshold, an even smaller percentage of the genome[20]. A survey of "never mutated" genes in the TCGA database, found only 5% of the genes fit into this category[22]. Even in normal germline cells, no more than 11% of mutations are projected to reduce organismal fitness[23], but the evolutionary constraints on tumor cells are expected to be far less than on multicellular organisms[24]. Further, Martincorena et al.[9] performed an extensive analysis of nonsynonymous/synonymous mutation ratios in somatic tissues, both cancerous and normal tissues. They concluded that only 0.02% to 0.5% of genes showed evidence of negative selection. They estimated that an average of only 0.5 mutations per consensus tumor genome were purified out by negative selection. Thus, only a minor fraction of diversity is expected to be removed by negative selection.

*Average mutation frequency calculated across subclones mutating at different frequency.*
The evidence so far accumulated indicates that the most prevalent types of mutations in cancer are single-base substitutions. This observation was probably unexpected, as these types of mutations are not associated with viral or environmental carcinogens. Viral agents usually insert or delete genomic sequences, while many chemical carcinogens act indirectly and form bulky adducts that are removed by nucleotide excision repair. Prominent in formation of single-base substitutions are errors in DNA replication resulting from misincorporation by DNA polymerases or diminutions by nucleotide-base repair. Major likely sources for single-base substitutions would be the major replicating enzymes, DNA polymerases –δ and –ε. These enzymes and interactive proteins are responsible for copying the three billion nucleotide base pairs in the human genome with only one or a very few mistakes. Errors by these enzymes or deficits in repair seem a likely cause of mutations as initially postulated in the concept of a mutator phenotype. If replicative DNA polymerases are unable to copy past bulky lesions in DNA, human cells have an armamentarium of error-prone DNA polymerases (Y – class) that can be brought to a stall to bypass the alteration in the DNA.

Data in this manuscript documents mutations in DNA polymerase genes, and, while most of these are not known to have functional consequences, it highlights the possibility that subclones may exist with differing mutations in the genomic maintenance machinery, resulting in differing rates of evolution. The mathematical treatment herein assumes that the effective mutation frequency is a population-weighted average of these different subclones and their mutation frequencies, and that this population weighted average is approximately constant at different levels of subclonal frequency.

We have observed a linear relationship between the logarithm of the fraction of apparently unmutated bases and sequencing depth. If serial mutator mutations progressively accumulated during tumor growth, and these subclones represented a large enough subset to alter the averages, one would expect subclones which formed later in tumor growth to have higher mutation rates and to generate more unique variants. Due to the branching nature of tumor evolution, these mutator subclones, progressively arising later, would correspond to rarer variants and the absolute value of the slope should increase with increasing depth, resulting in downward curvature (more negative slope), contrary to observation. Moreover, this downward curvature would produce linear regressions with positive y-intercepts, again contrary to observation. The data do not rule out the existence of hypermutator clones. However, there is no evidence from the data for hypermutator clones being prevalent enough to skew the averages computed by the model. A selective advantage for hypermutator clones may appear under selection pressures by therapy, which might favor subclones that can more rapidly acquire multiple resistance mutations[16]. Under those circumstances, these hypermutator clones may increase in relative prevalence or even become the dominant clone. However, the current study involves samples obtained at diagnosis, and under neutral evolution in the absence of therapy the proportion of hypermutator subclones might be expected to remain relatively constant. As long as the population weights of the different subclones are stable over time, $k_{mut-eff}$ can be calculated as a population weighted average.

*Extrapolation from the genes sequenced to the entire genome.*
Analyses discussed in the main article saw only minor differences in mutation frequency between the various genes sequenced. While there is known sequence variation in the mutation frequency throughout the genome, it seems most unlikely that all of the chosen genes with similar mutation frequencies were mutation hotspots. Therefore, it is more likely that they represent a reasonable approximation of an average mutation frequency.

*Extrapolation from the maximum depths achieved to every cell in the tumor.*
The estimate of total mutational burden of the tumor provided, and the conclusions about the likelihood of an arbitrarily selected site being mutated, depend on extrapolation of the depth achieved in this study to single cell depth representing every cell in the tumor. Although this study achieved unprecedented sequencing depth, we cannot rule out the possibility that the linear curves seen in Figure S1 might begin to level off (i.e., develop upward curvature) at still greater depth.

#### *Dependence on growth pattern and evaluation of consequences for estimation of actual and effective mutation rate in tumors[26])*

The results described above correspond to an average mutation frequency of 7.1 X $10^{-7}$ mutations per base per effective cell division. The actual number of cell divisions and average mutation frequency per base per actual cell division cannot uniquely be determined from this analysis, as they depend on the actual cell birth and death rates, whether these are constant, and whether the growth pattern is exponential or Gompertzian. Table S4 illustrates various scenarios and the numbers of actual cell divisions and mutation rates per actual cell division, which would be consistent with the data for the given scenario.

As illustrated in row 1 of the table, if there is simple exponential growth and no cell death, actual and effective mutation frequencies and actual and effective cell division numbers are identical. The birth and death rates from row 2 were estimated from colorectal cancer clinical data[26]. As illustrated in rows 2 and 3, as the birth and death rates become more closely matched, it takes more actual cell divisions to successfully increase the cell numbers by 1, and the actual number of cell divisions increases with a compensatory decrease in the average mutation frequency per actual cell division to account for the observed results.
In Gompertzian growth, the exponential growth rate exponentially decreases, reaching zero when the number of cells in the tumor reaches the carrying capacity C. At any given moment, the growth dynamics are given by the Gompertz differential equation:

$$\frac{dn}{dt} = g_0 n \frac{\ln(C) - \ln(n)}{\ln(C)}$$

(Equation S23),

where n is the number of cells in the tumor, C is the maximum carrying capacity, and $g_0$ is the initial net growth rate. We assume in the calculations and simulations for the table that the carrying capacity is $10^{10}$ cells, 10 $cm^3$, a conservative assumption, since larger lesions are commonly seen, and that the tumor grows to $10^9$ cells by the time of diagnosis.

Gompertzian growth dynamics can be implemented in several ways. In row 4, we make the assumption that birth and death rates both proportionally decrease with increasing tumor size until they both equal zero at the carrying capacity. This maintains a constant ratio of birth to death rates, and, therefore, equation 1 Methods can be integrated and equation S6 still holds. In this simplified case, Gompertzian growth dynamics is straightforward to analyze (row 4) and produces identical results to the comparable exponential case.

However, a more realistic Gompertzian implementation may be a constant death rate and the birth rate decreasing to equal it at the carrying capacity. In this case,

$$\frac{b-d}{b} = \frac{\ln(C) - \ln(n)}{\ln(C)}$$

(Equation S24),

where the actual number of cell divisions required to produce a new cell is b/(b-d). This case was simulated by dividing the growth from a single founder cell to $10^9$ cells into 30 doublings, with each doubling requiring more cell divisions, due both to the number of "effective" cell divisions needed to double the numbers, and to the increasing value of b/(b-d). The total number of cell divisions is given by:

$$Total\ number\ of\ cell\ divisions\ = \sum_{doublings, n=1}^{n=N/2} \frac{n\, b(n)}{b(n) - d(n)} = \sum_{doublings, n=1}^{n=N/2} \frac{n\ \ln(C)}{\ln(C) - \ln(n)}$$

(Equation S25)

It should be noted in Equation S25 that the estimate of total cell divisions is an n-weighted sum and is dominated by the larger values of n where more actual divisions are required to create a single new cell. In contrast, as shown above by equation S4, the observed mutations from each doubling are equal in number. This is because, even though later doublings involve more cells, mutations arising in any single cell are in rarer subclones and are less likely to be observed at limited depth. These effects cancel each other out. Thus, the total observed mutations are proportional to an unweighted sum of b/(b-d) = ln(C)/[ln(C) – ln(n)].

As a result of this effect, in row 6 for example, the number of actual cell divisions increases relative to row 1 by 7.2-fold, potentially implying that the average mutation frequency per actual cell division should be 7.2-fold lower than the effective mutation frequency derived from the analysis. However, in this scenario, a large fraction of the actual mutation burden is generated later in the process of tumor growth when the tumor begins to approach 10% of the carrying capacity and, therefore, there are many actual cell divisions per effective cell division. These later mutations are present only in very rare subclones since they arise very late in tumor development, and would go largely unobserved, even at the high depths achieved in this study. Hence, the effective mutation frequency is 3-fold underestimated in this scenario, and, thus, the actual mutation frequency, while 7.2-fold lower than the true effective mutation frequency, is only 2.4-fold lower than the apparent effective mutation frequency.

At high enough depth in the scenarios of rows 5 and 6, increasing mutations should be detectable corresponding to these rare subclones, leading to downward curvature (increasing negative slope) and an apparent positive intercept of the lines in Figures S1a-e. These were not observed, either because the Gompertzian models in rows 5 and 6 are not applicable or because the depth was insufficient to demonstrate this effect.
Row 5 utilizes the birth and death rates from Diaz et al.[26], as starting values when the tumor is small. Row 6 assumes a death rate of zero and an exponentially decreasing birth rate.

The effective mutation rate per nucleotide per new cell added to the tumor is the experimental observable in this work. The actual mutation rate per nucleotide per actual cell division cannot be determined without knowledge of the proliferation history of the tumor. For a given measured effective mutation rate, the inferred underlying actual mutation rate will be lower the more closely matched cellular birth and death rates are. When cell birth and death rates are closely matched, each effective cell division, i.e. increase in cell number by one, reflects more actual cell divisions.

The actual mutation rate per nucleotide per actual cell division reflects the underlying biophysical properties of the cell's DNA repair and replication machinery, and cannot change without a change in mutagens in the environment or a change in the DNA repair or replication machinery. We assume the actual mutation rate to be constant for much of tumor growth (see assumptions and approximations of model above).

In some growth patterns, such as Gompertzian growth, the cell birth and death rates become more closely matched as the tumor grows. Given a constant actual mutation rate, the effective mutation rate will thus increase over time. However, mutations occurring later in the tumor's history are rarer and harder to detect. Thus, sequencing fewer than every cell in the tumor will, under Gompertzian growth, lead preferentially to detection of mutations from earlier timepoints in the tumor's history, prior to the increase in effective mutation rate. This will lead to an underestimate of the average effective mutation rate over time, and an underestimate of the mutational burden within the tumor. Our conclusion that no nucleotide locus is wild type in every cell will be even more definite under these circumstances.

We have observed, as predicted for exponential growth, a straight line of the curve of the natural logarithm of the unmutated fraction of sites when plotted against sequencing depth. Gompertzian growth predicts increasing slope, and therefore curvature, as the effective mutation rate continually increases. Thus, we find no evidence for Gompertzian growth as far forward as we can see in time. We cannot rule out that Gompertzian growth may occur later in tumor development; this would only further increase the mutation burden as described above.

*Consequences for the mutator hypothesis*[23,27]
The mutator hypothesis states that the actual mutation frequency per base per cell division in tumors is higher than the actual mutation frequency per base per cell division in normal tissues. Lynch (2010) reviews a variety of estimates of mutation frequencies in normal tissues[23], including studies of the development of retinoblastoma in heterozygous carriers, studies of the APC gene in intestinal epithelia, studies of the HGPRT genes in cultured lymphocytes and fibroblasts, and studies of the PigA gene in cultured lymphocytes. These estimates range from $2.7 \times 10^{-10}$ to $1.5 \times 10^{-9}$ mutations per base per actual cell division. Roach et al. sequenced whole genomes of a family of four and only 70 mutations were identified between generations[27]. They estimated a human inter-generational mutation frequency of $1.1 \times 10^{-8}$ per nucleotide position per haploid genome. Assuming that by adulthood an individual gamete undergoes 100 cell generations, the mutation frequency per cell generation would be $1 \times 10^{-10}$ per nucleotide.

In contrast, the estimated mutation frequencies per single base per actual cell division in colorectal tumors in a large variety of tumor growth scenarios including Gompertzian growth are illustrated in Table S4, ranging from $2.8 \times 10^{-8}$ to $7.1 \times 10^{-7}$, a numerically higher range than estimated for normal tissues. This comparison would appear to generally support the mutator hypothesis. However, this conclusion cannot be definitive as the actual growth patterns of both normal tissues and tumors are still unknown.

**Probability of existence of at least one tumor cell with a mutation at a given base.**
Mathematical models of mutational acquisition of drug resistance in cancer have been created by many groups beginning with Goldie and Coldman[28]. Based on this framework, other modelers[29,30] concluded that drug resistance to single agent therapy might frequently be pre-existing. Loeb et al[31] and Sottoriva et al[12] pointed out that mutational diversity leading to drug resistance would be further enhanced in the

presence of a mutator phenotype, while Sottoriva and colleagues[5,12] emphasized the point that neutral evolution would also enhance diversity.

Bozic et al[7] modeled the acquisition of drug resistance using a branching pathways approach, demonstrating in detail that single agent therapy would inevitably lead to drug resistance, that the probability of pre-existing resistance would continuously increase with tumor size, and that simultaneous combinations would outperform successive single agents if the components of the combination could be safely given at effective dosages. Beckman, Schemmann, and Yeang[16] and Yeang and Beckman[32] studied this problem in the setting where simultaneous combinations require meaningful dose reduction due to toxicity. They found that frequent adaptation with pulses of simultaneous combinations and full dose monotherapy governed by an evolutionary model and subclonal tracking greatly outperformed the application of simultaneous combinations until failure.

The above authors all discussed the *likelihood* that *many* instances of drug resistance are due to pre-existing resistant subclones at diagnosis, and this idea has been validated in numerous studies where resistance mutations found at clinical relapse were detected as minority subclones at diagnosis[33]. However, these studies relied on low depth sequencing. Moreover, many of these approaches relied on the infinite sites assumption, leading to underestimates of the total mutational burden in tumors sufficiently large to be clinically detected, as discussed above.-Based in the current work we conclude that pre-existing resistance to single agent therapy is *universal* and *inevitable.*

The average number of cells harboring any mutation of interest will be $k_{mut\text{-}eff}$ N, which for a tumor of $10^9$ cells means there will be on average 700 cells resistant to any single agent at diagnosis, even if there is only one mutational mode of resistance. Moreover, we can calculate the likelihood of resistance to more than one non-cross resistant therapy occurring simultaneously in the same cell, finding it is low at diagnosis but that it increases as the tumor grows, in agreement with Bozic et al[7]. However, given our highly accurate sequencing methods, and our theoretical model which is free of the infinite sites assumption and therefore applicable to clinically relevant tumor sizes, our estimates of these quantities will be more accurate and more generally applicable. These likelihoods correspond only to resistance due to mutations alone. As there are other sources of resistance (see below), we can at times see immediate resistance to non-cross resistant combinations in the clinic.

From the average value of $k_{mut\text{-}eff}$ of 7.1 X $10^{-7}$, we determine the likelihood that every cell is unmutated at any given neutral single base locus at diagnosis. We assume the tumor burden ranges from 1 $cm^3$, the approximate limit of computed tomography detection, to 10 $cm^3$, representing multiple or slightly larger lesions at diagnosis. Many authors assume $10^9$ cells/$cm^3$, but a recent study[14] suggests there could be as few as $10^8$ cells/$cm^3$. Thus, depending on the size of the tumor at diagnosis and their cellular density, a patient could have $10^8 - 10^{10}$ malignant cells. Assuming $10^9$ cells, and applying equation S8, the probability that any given locus is unmutated at every cell in such a small lesion is $10^{-308}$. Thus, it is extremely unlikely that there is any nucleotide in DNA that is wild type in every cell, even at diagnosis**.** This result is robust. If we have overestimated the effective mutation frequency by a factor of 10, the probability of a neutral single base locus being unmutated in every cell in the lesion is $10^{-31}$. For the probability of a neutral single base locus being unmutated in every cell in the tumor being as high as 1%, we would have had to have overestimated the effective mutation frequency by 150-fold. The above figures become even more striking as a cancer grows from $10^9$ cells at diagnosis to $10^{11}$-$10^{12}$ cells in the terminal phase.

It is apparent from the above discussion that any given genomic position that is capable of encoding resistance to any therapy is highly likely to be mutated in at least one tumor cell at diagnosis, unless mutation of the position confers a significant fitness disadvantage in the absence of therapy. Moreover, it is likely in most cases that multiple single base loci in the genome confer resistance; further increasing the likelihood that at least one resistant cell will be present. While the number of single base sites in the genome conferring a resistance phenotype is unknown, and likely varies depending on the therapy, a common assumption is that there are on the order of 100 sites in the genome, mutation of which may confer resistance[18,26].

These considerations raise the obvious point of the need for more than one non-cross resistant component of therapy, whether given as simultaneous combinations[7] or as a complex sequence of simultaneous combinations and monotherapy pulses[16,32]. Assuming complete non-cross resistance, and independence of acquisition of the resistance mutations, the probability $\mathbf{P}_{\textbf{no simultaneous resistance}}$, of having no single cell simultaneously resistant to **K** non-cross resistant agents, for which each has **R** sites in the genome, mutation of which can cause therapy resistance, in a tumor of N cells is approximately, -in analogy with equation S8:

$$P_{no\ simultaneous\ cross\ resistance} = e^{-(R\, k_{mut-eff})^{K} N}$$

(Equation S26)

In table S5, we calculate values of $P_{\text{no simultaneous resistance}}$ for a number of scenarios of treatment with 2, 3, and 4 non-cross resistant therapies as a function of tumor burden N ($10^9$-$10^{12}$) and number of single base sites in the genome conferring resistance (R), assuming $k_{mut-eff} = 7.1 \times 10^{-7}$

Several generalizations are evident from the table. One is that the required number of non-cross resistant agents to ensure no fully resistant cell depends heavily on both the tumor burden and the number of bases in the genome that confer neutral fitness resistance mutations. The table suggests that, in the event of a single base in the entire genome that confers resistance, 2 non-cross resistant therapies are required for low-moderately low tumor burdens ($10^9$-$10^{10}$ cells), whereas 3 non-cross resistant therapies are required for higher tumor burdens in the $10^{11}$-$10^{12}$ range. However, if 100 neutral single base loci confer resistance to each agent, the number of required non-cross resistant agents rises to 3 and 4 in low/moderate and high tumor burdens respectively.

Yet, the challenge is even greater for a variety of reasons[3449]:

- The above does not consider other genetic mechanisms of resistance such as chromosomal rearrangements and copy number variation (CNV). CNV also increases the number of alleles available for dominant mutations, while making homozygosity more unlikely in the case of recessive mutations.
- The above does not consider epigenetic changes leading to resistance.
- The above does not consider non-genetic forms of resistance, such as feedback loops hard-wired into cells. Each non-cross resistant therapy may itself consist of several individual agents designed to hit several nodes of a signalling network corresponding to only one genetic state.
- The above does not account for partial effectiveness of therapy, inhomogeneous distribution of therapies into tumor tissues, or interactions of tumor cells with each other or with the host tissues.

**Simulations of unique subclones observed as a function of sequencing depth in the presence and absence of selection: sensitivity for ruling out selection model.**
Exact simulations for these cases were performed using deterministic models.

For the neutral evolution case, we use equation S6 above for the fraction of unmutated bases, a capture set of 10 kilobases, and $k_{mut-actual}$ of $2.0 \times 10^{-7}$ per nucleotide per actual cell division, which fits our data given the birth and death rates of 0.25/day and 0.18/day estimated based on experimental data[17] (line 2, Table

S4). The number of mutated sites is calculated from the fraction unmutated at the depths indicated in Figure 2 (main text).

In the selection case, we consider three groups of loci, i = 1, 2, 3 where i = 1 is positively selected loci, i = 2 is neutral loci, and i = 3 is negatively selected loci.

Let s equal the relative advantage of each group over its nearest inferior category. Let $r_i$ denote the fitness of group i. We assume $r_2 = 1$, $r_1 = 1 + s$, $r_3 = 1 - s$ in the simulation although the equations below are more general.

Let b be the common birth rate for all these cells, and let $d_i$ be adjusted death rates calculated from the relative fitnesses assumed above. Further, let the age of the tumor from the formation of the founder cell to the time of biopsy be T in days, or bT in cell generations, where b is the birth rate in cell generations/day. Cells formed at time t with r > 1 are enriched, and cells with r < 1 depleted, according to:

$$Enrichment\ factor = e^{(r_i-1)b(T-t)}$$

(Equation S27)

For any of the three types of mutations, the number of detectable mutations at depth D is the time integral of the instantaneous number of detectable mutations at time t from 0 to T. As demonstrated above, the probability of detecting a neutral mutation in a particular locus from a cell formed at time t is a constant, $k_{mut\text{-}eff}$ D, and therefore may be taken outside of any integral over time. However, for positively and negatively selected genes, this constant is multiplied by an enrichment factor that is not constant (equation S27), and when integrated over time from 0 to T yields an average enrichment factor:

$$Average\ enrichment\ factor = f(r_i, T) = e^{(r_i-1)b\,T}\ \frac{\int_0^T e^{-(r_i-1)\,bt}\,dt}{T} = \frac{e^{(r_i-1)bT} - 1}{(r_i - 1)bT}$$

(Equation S28)

As a check on this, we can show that the limit of the enrichment factor as $r_i \rightarrow 1$, or as $T \rightarrow 0$ is 1, as would be expected in the limit of no enrichment (expand the exponential in a Taylor series and truncate at the linear term, or apply L'Hôpital's rule directly to obtain these limits).

With this enrichment factor, and the fact that D and $k_{mut\text{-}eff}$ are constants, we observe that:

$$Expected\ number\ of\ observed\ mutations\ per\ nucleotide\ locus\ of\ type\ i = k_{mut-eff} f(r_i, T) D$$

(Equation S29)

Let us further denote the relative prevalence of the three type of sites by $a_i$, where $\sum_{i=1}^{3} a_i = 1$. Finally, for convenience, let us term $\beta_i = k_{mut\text{-}eff}\ f(r_i, T)$.

Then, the fraction of unmutated sites of type i is

$$F_{apparent-unmutated_i} = e^{-\beta_i D}$$

(Equation S30)

and the fraction of unmutated sites overall is

$$F_{apparent-unmutated\ with\ selection} = \sum_{i=1}^{3} a_i\ e^{-\beta_i D}$$

(Equation S31).

The simulations utilize T = 428 days representing 30 doublings from founder cell formation given the net doubling rate of 0.07 (b-d). Thirty doublings is sufficient for the single founder cell to grow to a diagnosable lesion with $10^9$ cells.

For the selection coefficient s, we have utilized values from 0.04 to 0.25. Bozic et al[25] have modeled the selection coefficient based on evaluation of putative passenger and driver mutations in astrocytic glioblastoma and pancreatic adenocarcinoma sequences, and validated their conclusions by predicting the kinetics of appearance and growth of polyps in familial adenomatous polyposis in two of three datasets. They reached the conclusion that, even for APC, s is very small: 0.004. However, the key equation in their analysis is equation 2, in which s appears in a ratio with the mutation rate. They then use a mutation rate three orders of magnitude lower than what we have determined. Using our mutation rate, their equation 2 would give a value of s of nearly 4, which is in accord with the value of 2.76 more recently measured by direct observation of colonic stem cell crypt evolution using fluorescently labeled cells in genetically engineered mice[35]. Moreover, since the modeling of polyp appearance kinetics does not account for the likelihood of elimination of most nascent polyps by immune surveillance and other factors which may confound comparisons of tumor initiation kinetics with clinical observations[10], the polyp modelling may not further determine their parameters. Williams et al[36] have examined bulk sequencing data from multiple sources, looking for deviations from neutral evolution in the curve of variant allele frequency versus total mutation burden (as a surrogate for time). They have not detected evidence of a value of s below 0.2, which they also state as their limit of sensitivity.

We varied $a_1$ from 0.004 to 0.1 and s from 0.04 to 0.25, keeping $a_1 = a_3$ for convenience (the value of $a_3$ does not matter as it is unlikely we have sequenced deeply enough to find mutations at negatively selected sites). For each combination of $a_1$ and s, we varied $k_{mut-eff}$ and the y intercept to obtain the best fit to experimental data, plotted as ln ($f_{apparent-unmutated}$) versus depth. Based on the various population genetic and experimental estimates for normal tissue discussed above, $k_{mut-actual}$ must be $\geq 10^{-10}$, and since $k_{mut-eff} \geq k_{mut-actual}$, we also have $k_{mut-eff} \geq 10^{-10}$. For both neutral and selection models, best fit absolute residuals were evaluated as a fraction of their corresponding datapoint, resulting in an average fractional absolute residual and a 95% confidence interval thereof for both the selection model and the neutral model. Selection models were considered ruled out when their average fractional absolute residual exceeded the upper 95% confidence limit of the comparable statistic for the neutral model. In Figure S2, we see that for $a_1 = 0.015$, selection models with $s \geq 0.23$ are ruled out.

We thus found, in agreement with Williams et al[36], that we could rule out selection of $s \approx 0.2$ or stronger, as optimal fit to the data required $k_{mut-actual} \leq 10^{-10}$. We note[35] that values of s for strong drivers range from about 1 to 4. It is difficult to rule out selection if $a_1 < 0.01$ as it determines the amplitude of curvature.

The ability to rule out selection models improves if we demand that the graph intersect the origin, a theoretical point corresponding to no unique subclones observed at a sequencing depth of zero. Further, if we demand that $k_{mut-eff}$ be that estimated from the linear model, the selection model is more easily ruled out. Figure 2 in the main text demonstrates this in that when $k_{mut-eff}$ is constrained to be the same as the neutral model, a selection model with $a_1 = 0.015$ and $s = 0.125$ is easily ruled out by visual inspection. Figure 2 in the main text is graphed according to the more intuitive Williams et al[5] presentation, an approximation that is valid for the sequencing depths utilized in this study.

In principle, exact simulation of the selection model will in general not give a straight line, but a triphasic curve with lesser absolute slope with increasing duplex sequencing depth, resulting in curvature (upward slope with downward curvature for the Williams et al[5] formulation; downward slope with upward curvature for our approach). The initial phase at low depth (high variant fractions) is dominated by selected loci which accumulate rapidly in the majority of cells. Then a second phase follows representing the neutral loci which accumulate more slowly and at lower variant fractions, and thus become visible at higher depth. Finally, a third phase representing small numbers of negatively selected genes at still lower variant fractions may be observed. If we had modeled a continuous distribution of fitness, there would be a continuous downward curvature rather than 3 phases. However, the ability to observe all three phases, or indeed to observe curvature, depends on the parameters and the sequencing depths chosen. We can't rule out very weak selection or selection occurring at a very small fraction of loci. However, these latter two selection cases are themselves close to neutral evolution.

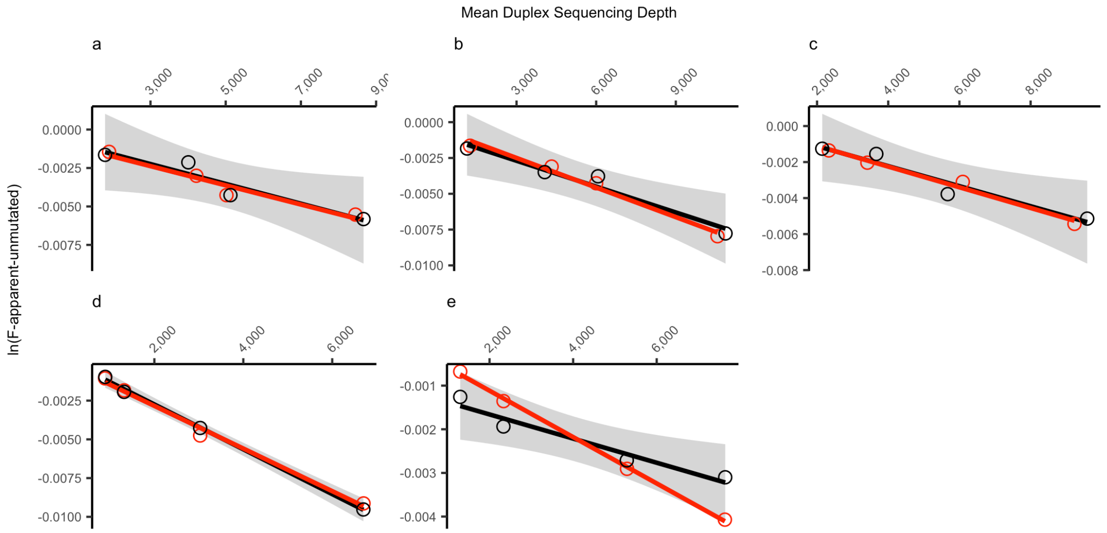


**Fig. S1.** ln ($F_{apparent\ unmutated}$) versus sequencing depth for tumors 1 – 5 (a – e), Table S3. Black symbols represent experimental data. Red symbols represent subsampled data. Line represents regression line through points shown. Shaded area is the 2-sided 95% confidence interval of the experimental data as determined by the student's t test with 2 degrees of freedom. Of the total of 20 instantiations of subsampling, 19/20 (95%) are within the 95% confidence interval of the experimental data. The exception is shown in sub-figure e.

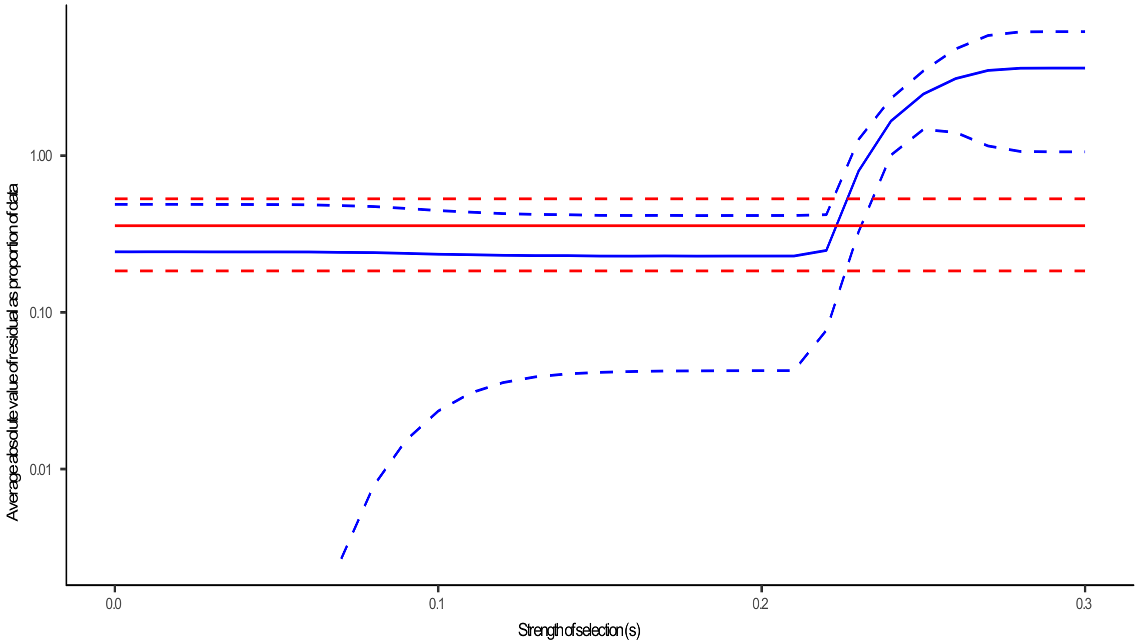


**Fig. S2.** Sensitivity threshold for ruling out selection models. Average absolute fractional residual curves for best fit neutral (red) and selection (blue) models are plotted (solid lines) along with their 95% confidence limits (dashed lines) as a function of the strength of selection s. Reference data was one of the 5 fresh frozen CRC tumors that we sequenced at multiple depths, plotted according to the approach in Online Methods, with the natural logarithm of the unmutated fraction of the capture set plotted against depth. For the selection model, $a_1$ was set at 0.015, indicating 1.5% of loci were positively selected (97% were neutral and 1.5% negatively selected). For each model $k_{mut\text{-}eff}$ and the y intercept value (theoretically zero) were varied in a search for the optimal fit. $k_{mut\text{-}eff}$ was constrained to be $\geq 10^{-10}$. For each point in the data, absolute values of the residual as a fraction of the data point itself were recorded. These values were averaged across the data points for each model. For $s > 0.23$ the average absolute fractional residual is greater than the upper 95% confidence limit of the same statistic for the neutral model, suggesting that this strength of selection is ruled out. Strong drivers typically have s on the order of 1-4[50]. See Supplemental Methods for details of simulation.

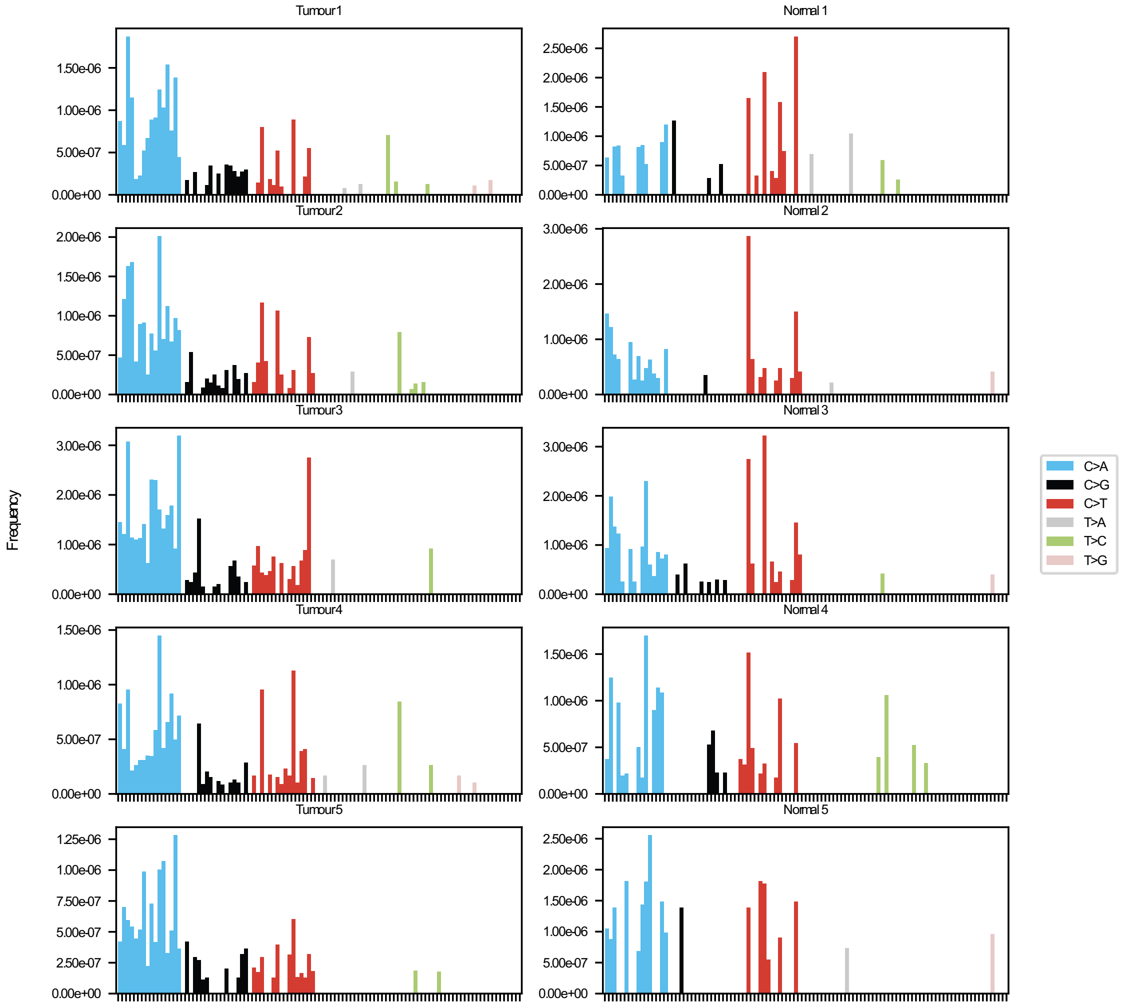


**Fig. S3.** Signatures of mutation for tumors and associated normal samples used for regression analysis. Frequencies represent counts of mutations divided by total number of instances of that context sequenced.

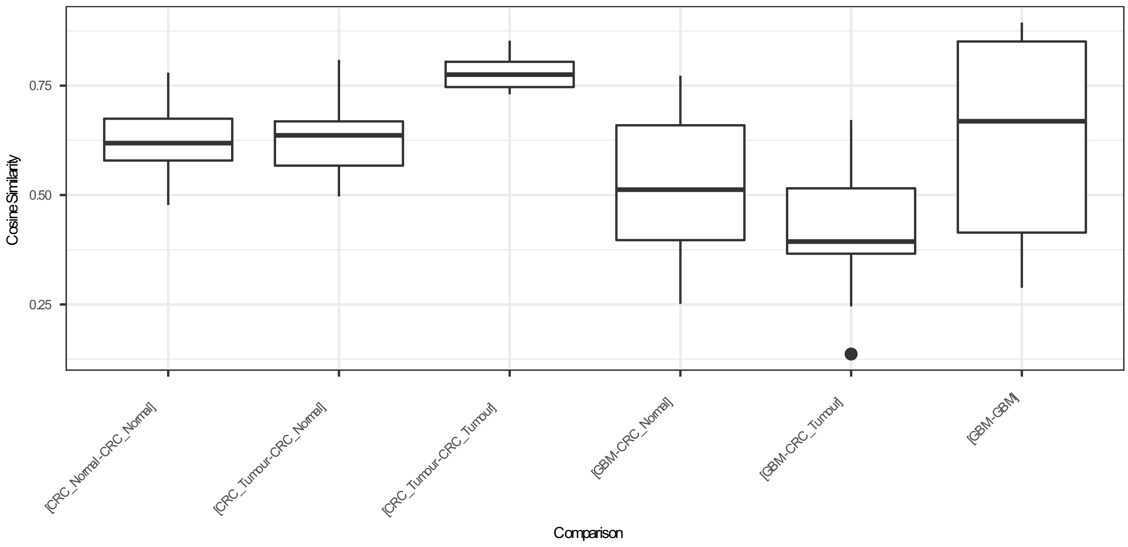


**Fig. S4.** Plot of cosine similarities of mutation signatures, grouped by type of comparison. These results show that, from the context of a tumor sample, there is a significant difference between another tumor sample and a GBM sample. Detailed results of an anova of this data are discussed in Table S6.

**Table S1. Oligonucleotide sequences used. ID corresponds to Truseq indexes; N = random nucleotide sequence; I = inosine.**

| ID | Sequence |
|---|---|
| MWS13 | AATGATACGGCGACCACCGAG |
| MWS20 | GTGACTGGAGTTCAGACGTGTGC |
| MWS21 | CAAGCAGAAGACGGCATACGAGATXXXXXXGTGACTGGAGTTCAGACGTGTGC |
| MWS51 | AATGATACGGCGACCACCGAGATCTACACTCTTTCCCTACACGACGCTCTTCCGATCT |
| MWS55 | TCTTCTACAGTCANNNNNNNNNNNNAGATCGGAAGAGCACACGTCTGAACTCCAGTCAC |
| MWS60 | AATGATACGGCGACCACCGAGATCTACACTCTTTCCCTACACGACGCTCTTCCGATCTIIIIIIIIIIIITGACT |
| MWS61 | GTCAIIIIIIIIIIIIAGATCGGAAGAGCACACGTCTGAACTCCAGTCAC |

**Table S2. Parameters used in the core model or in the discussion.** Input parameters are experimentally measured or inferred (number of cells in a tumor inferred from size). Output parameters are derived from the analysis. Several parameters not used in the model are discussed herein to clarify their relationship to the model. *Number of effective cell divisions is the net number of events resulting in a net increase in the tumor population by 1 cell. For a tumor of N cells originating from a single founder cell, the number of effective cell divisions is by definition N-1. Hence the parameter $N_E$ is directly estimable from the tumor size. **Subclonal mutations are called relative to the tumor clonal reference sequence, not the germline sequence. The tumor clonal reference sequence is presumably derived from the founder cell(s).

| **Parameter** | **Symbol** | **Use in current model** |
|---|---|---|
| Number of effective cell divisions* | $N_E$ | Derived output parameter |
| Number of actual cell divisions | $N_A$ | Not used in core model |
| Cell birth rate | b | Not used in core model |
| Cell death rate | d | Not used in core model |
| Number of cells in tumor at time of analysis | N | Input parameter |
| Number of cells in tumor at earlier time t | n(t) | Intermediate parameter used in mathematical formalism |
| Mutation frequency per base per effective cell division* | $k_{mut\text{-}eff}$ | Derived output parameter |
| Mutation frequency per base per actual cell division | $k_{mut\text{-}actual}$ | Not used in core model |
| Duplex Sequencing depth | D | Input parameter |
| Fraction of bases sequenced at which no unique subclonal mutation is observed** | $F_{apparent\text{-}unmutated}$ | Input parameter |

**Table S3. Range of depths, number of independent depths sequenced, linear correlation coefficient R, square of the linear correlation coefficient (fraction of variation of data explained by linear model), estimated $k_{mut\text{-}eff}$ (minus the slope), y-intercept, and its two-sided 95% confidence interval based on t-test with 2 degrees of freedom for 5 colorectal tumors plotted according to equation 8. Results shown graphically in Figures S1a-e.**

| Tumor | Range of depth | Number of points | R | $R^2$ | $k_{mut\text{-}eff}$ | y-intercept | 95% confidence interval, y-intercept |
|---|---|---|---|---|---|---|---|
| 1 | 1792-8665 | 4 | 0.953 | 0.908 | $6.44 \times 10^{-7}$ | $-3.12 \times 10^{-4}$ | $+3.12$ to $-3.74 \times 10^{-3}$ |
| 2 | 1137-10872 | 4 | 0.979 | 0.958 | $6.02 \times 10^{-7}$ | $-8.89 \times 10^{-4}$ | $+1.62$ to $-3.39 \times 10^{-3}$ |
| 3 | 2146-9590 | 4 | 0.967 | 0.935 | $5.56 \times 10^{-7}$ | $-2.21 \times 10^{-6}$ | $+2.66$ to $-2.67 \times 10^{-3}$ |
| 4 | 895-6700 | 4 | 0.999 | 0.999 | $1.45 \times 10^{-6}$ | $1.46 \times 10^{-4}$ | $+7.83$ to $-4.90 \times 10^{-4}$ |
| 5 | 1299-7639 | 4 | 0.972 | 0.944 | $2.77 \times 10^{-7}$ | $-1.11 \times 10^{-3}$ | $-2.09 \times 10^{-4}$ to $-2.09 \times 10^{-3}$ |

**Table S4. Effective and actual cell division numbers and mutation frequencies per base per effective or actual cell division for different growth scenarios, including varying cell birth and death rates, exponential growth, and several variations of Gompertzian growth. [1]Calculated using equations 1 and 8; [2]Birth and death rates as published[8]; [3] Calculated using equations 1 and 8. [4]Simulated using equations 1, 8, 12, and 13, assuming the carrying capacity (defined below) is $10^{10}$ cells and the tumor is sampled at a size of $10^9$. [5]In the scenarios in the bottom two rows, the effective mutation frequency is an *apparent* effective mutation frequency. The true effective mutation frequency is approximately three fold higher as determined by the simulation.**

| | Model | Growth pattern | Birth rate (b) (per day) | Death rate (d) (per day) | Number of actual cell divisions (founder cell to $10^9$) | Net number of effective cell divisions (founder cell to $10^9$) | Actual mutation frequency (per base per actual cell division) | Effective mutation frequency (per base per effective cell division)[5] |
|---|---|---|---|---|---|---|---|---|
| 1 | Exponential[1] | | 0.25 | 0 | $10^9$ | $10^9$ | 7.1 X $10^{-7}$ (per base per actual cell division) | 7.1 X $10^{-7}$ |
| 2 | Exponential[1,2] | | 0.25[2] | 0.18[2] | 3.57 X $10^9$ | $10^9$ | 2.0 X $10^{-7}$ | 7.1 X $10^{-7}$ |
| 3 | Exponential[1] | | 0.25 | 0.24 | 2.5 X $10^{10}$ | $10^9$ | 2.8 X $10^{-8}$ | 7.1 X $10^{-7}$ |
| 4 | Simplified Gompertzian[2,3] | b and d decreasing in proportion with increasing tumor size | 0.25, decreasing to 0 at carrying capacity | 0.18, decreasing to 0 at carrying capacity | 3.57 X $10^9$ | $10^9$ | 2.0 X $10^{-7}$ | 7.1 X $10^{-7}$ |
| 5 | Gompertzian[2,4] | Relative survival probability (b-d)/b decreasing with increasing tumor size | 0.25, decreasing to 0.18 at carrying capacity | 0.18 | 2.57 X $10^{10}$ | $10^9$ | 8.4 X $10^{-8}$ | 7.1 X $10^{-7}$ |
| 6 | Gompertzian[2,4] | Relative survival probability (b-d)/b decreasing with increasing tumor size | 0.25, decreasing to 0 at carrying capacity | 0 | 7.20 X $10^9$ | $10^9$ | 2.9 X $10^{-7}$ | 7.1 X $10^{-7}$ |

**Table S2. Probability $P_{no\ simultaneous\ cross\ resistance}$ that no cell in a cancer of N total cells will be resistant to all of K non-cross resistant therapies, where in each case there are R neutral single bases in the genome, mutation of which confers resistance.**

| Number of cells (N) | Number of neutral single base resistance loci (R) | Number of non-cross resistant therapies (K) | Probability that no cell in the tumor will be resistant to all therapies ($P_{no\ simultaneous\ cross\text{-}resistance}$) |
|---|---|---|---|
| $10^9$ | 1 | 2 | 0.999 |
| $10^9$ | 100 | 2 | $6.5 \times 10^{-3}$ |
| $10^9$ | 100 | 3 | > 0.999 |
| $10^{10}$ | 1 | 2 | 0.995 |
| $10^{10}$ | 100 | 2 | $1.3 \times 10^{-22}$ |
| $10^{10}$ | 100 | 3 | 0.996 |
| $10^{11}$ | 1 | 2 | 0.951 |
| $10^{11}$ | 1 | 3 | 1.000 |
| $10^{11}$ | 100 | 2 | $< 1 \times 10^{-100}$ |
| $10^{11}$ | 100 | 3 | 0.965 |
| $10^{11}$ | 100 | 4 | > 0.999 |
| $10^{12}$ | 1 | 2 | 0.604 |
| $10^{12}$ | 1 | 3 | > 0.999 |
| $10^{12}$ | 100 | 3 | 0.699 |
| $10^{12}$ | 100 | 4 | > 0.999 |

**Table S6. Range and p-values for Tukey HSD ad hoc test based on anova for data in Figure S4. GBM signatures are significantly different from those of CRC tumor or surrounding normal tissue. Regarding the comparison of CRC and surrounding normal, the uncorrected p value suggests a significant difference, but corrected for the multiple statistical comparison the p value is no longer significant.**

| Comparison | Mean Diff. | Lower | Upper | p value |
|---|---|---|---|---|
| [GBM-GBM] – [GBM-CRC_Normal] | 0.1079 | -0.0351 | 0.2509 | 0.2502 |
| [GBM-CRC_Normal] – [CRC_Normal-CRC_Normal] | -0.0939 | -0.2369 | 0.0491 | 0.4028 |
| [CRC_Tumor-CRC_Normal] – [CRC_Normal-CRC_Normal] | 0.0098 | -0.1332 | 0.1527 | 0.9999 |
| [CRC_Tumor-CRC_Tumor] – [CRC_Tumor-CRC_Normal] | 0.1508 | 0.0078 | 0.2937 | 0.0325 |
| [GBM-CRC_Tumor] – [CRC_Tumor-CRC_Tumor] | -0.3474 | -0.4904 | -0.2044 | 0.0000 |
| [GBM-GBM] – [GBM-CRC_Tumor] | 0.2009 | 0.0579 | 0.3438 | 0.0012 |

**Dataset S1 (separate file)**

Capture probes used for targeted capture in this study
